# PACMON: Pathway-guided Multi-Omics data integration for interpreting large-scale perturbation screens

**DOI:** 10.64898/2026.03.20.713295

**Authors:** Arber Qoku, Tyra Stickel, Sareh AmeriFar, Thomas Oellerich, Sebastian Wolf, Florian Buettner

## Abstract

High-throughput perturbation screens coupled with single-cell molecular profiling enable systematic interrogation of gene function, yet interpreting the resulting data in terms of biological pathways remains challenging. Existing approaches either identify latent gene modules without linking them to perturbations, or model perturbation effects without incorporating prior biological knowledge, limiting interpretability and scalability. Here, we introduce PACMON (Pathway-guided Multi-Omics data integration for interpreting large-scale perturbation screens), a Bayesian latent factor model that jointly infers pathway-level programs and their modulation by experimental perturbations. PACMON decomposes multimodal molecular measurements into shared latent factors aligned with known biological pathways through structured sparsity priors, while simultaneously estimating how each perturbation activates or represses these pathway programs. The framework naturally accommodates multiple data modalities and employs stochastic variational inference for scalable application to large datasets. We evaluate PACMON in three settings of increasing complexity. On synthetic data with known ground truth, PACMON achieves near-perfect recovery of pathway structure and perturbation effects, outperforming existing methods in both accuracy and computational scalability. Applied to a multimodal Perturb-CITE-seq screen of melanoma cells, PACMON recovers coherent interferon-signaling and cell-cycle programs spanning RNA and surface-protein modalities and identifies interpretable perturbation–pathway associations consistent with known immune-evasion mechanisms. Finally, we apply PACMON to the Tahoe-100M perturbation atlas — approximately 100 million cells and over 1,000 drug–dose combinations — producing the first pathway-level latent factor analysis at this scale and revealing biologically meaningful drug-response landscapes across Hallmark pathway programs. PACMON provides a unified, scalable and interpretable framework for mapping perturbation effects onto biological pathways in modern large-scale perturbation experiments.

## 1 Introduction

High-throughput perturbation experiments have become a cornerstone of modern functional genomics. Complementing single-cell readouts with targeted genetic or chemical interventions enables the systematic interrogation of how molecular systems respond to controlled perturbations [Dixit et al., 2016, Datlinger et al., 2017, Replogle et al., 2022]. Recent technical advances have further expanded the scale and scope of such experiments, making it possible to profile millions of cells across hundreds to thousands of perturbations in a single study, and increasingly across multiple molecular modalities [Mimitou et al., 2019, Norman et al., 2019, Zhang et al., 2025].

Interpreting these data in a biologically meaningful way remains challenging. Cellular regulatory systems are organised in a modular manner, with biological functions carried out by coordinated gene programs rather than isolated genes [Hartwell et al., 1999, Segal et al., 2003]. At the same time, perturbation responses are often sparse and modular, with most perturbations influencing only a limited number of co-regulated gene programs rather than inducing widespread, unstructured changes across the whole transcriptome [Yao et al., 2023, Zhou et al., 2023]. Despite this structure, perturbation analyses are often performed at the level of individual genes, typically yielding long lists of affected genes that are difficult to interpret while ignoring coordinated responses across gene modules. In addition, noise, sparsity and limited numbers of cells per perturbation reduce statistical power in single-cell settings [Finak et al., 2015, Srivatsan et al., 2020, Replogle et al., 2022]. Latent variable models provide a principled framework for addressing this challenge by representing high-dimensional molecular measurements in terms of a smaller number of latent factors that capture shared structure across genes or proteins and can be interpreted as gene modules [Stegle et al., 2015, Buettner et al., 2015, 2017, Qoku et al., 2025].

Recent work has extended this framework by explicitly linking perturbations to latent modules. For example, Factorize-Recover for Perturb-seq analysis (FR-Perturb) first identifies modules via a sparse PCA and, in a second step, links modules to perturbations via LASSO [Yao et al., 2024]. Guided sparse factor analysis (GSFA) consolidates these individual steps into a single framework and jointly infers factors and the effects of individual perturbations on modules [Zhou et al., 2023]. While these advances better account for the intrinsic correlation structure of the transcriptome, they still have important limitations. First, interpretation of inferred latent factors can be cumbersome, since they require careful analysis of factor weights and post-hoc annotation to pathways via enrichment tools [Reimand et al., 2019]. Second, these approaches are restricted to transcriptomic data and do not naturally accommodate multimodal perturbation assays. Finally, GSFA relies on Markov chain Monte Carlo (MCMC) for model inference, which prohibits application to modern large-scale perturbation screens due to poor computational scalability and prolonged runtimes.

In parallel, advances in interpretable factor analysis have demonstrated that incorporating prior biological knowledge directly into the model can yield latent factors that are interpretable by design, reducing the need for post-hoc explainability analysis of factor weights. For example, structured sparsity priors, strong regularisation and network based approaches have been introduced to explicitly align individual latent factors with specific pathways [Qoku and Buettner, 2023, Qoku et al., 2025, Kunes et al., 2024, Lotfollahi et al., 2023b]. However, these approaches are unsupervised and do not model perturbations, limiting their ability to directly quantify pathway-level perturbation effects.

Here, we introduce PACMON, a Bayesian latent factor model designed to infer associations between pathways and perturbations from large-scale perturbation screens. PACMON represents molecular measurements through latent factors linked to specific biological pathways and models these factors as functions of experimental perturbations. By combining perturbation-aware latent modelling with pathway-informed structured sparsity, PACMON directly quantifies how perturbations modulate a small number of biological programs rather than individual genes. The framework naturally extends to multimodal data and employs variational inference, enabling scalable application across a wide range of perturbation experiments.

## 2 Results

We evaluated PACMON in three increasingly complex settings. First, using synthetic perturbation data with known ground truth, we assessed recovery of pathways, perturbation effects and downstream differential-expression signals, while benchmarking scalability against existing methods (Fig. 2). Second, we applied PACMON to a multimodal Perturb-CITE-seq melanoma screen and showed that it recovers coherent pathway programs and interpretable perturbation-pathway associations across RNA and surface proteins (Fig. 3). Third, we analysed the Tahoe-100M perturbation atlas to demonstrate that PACMON scales to atlas-level data while preserving pathway-level interpretability and revealing biologically meaningful drug-response structures (Fig. 4).

### 2.1 PACMON provides an interpretable and scalable framework for multimodal perturbation analysis

PACMON is a Bayesian latent factor model that integrates multimodal molecular measurements with prior biological knowledge to infer interpretable perturbation–pathway associations from large-scale perturbation screens (Fig. 1). The model takes as input high-dimensional molecular profiles from one or more data modalities (e.g., RNA expression and surface-protein abundance) together with a perturbation design matrix encoding the experimental condition of each cell. These inputs are decomposed into three core components: (i) a shared set of latent factors representing pathway-level gene programs, (ii) modality-specific loading matrices that map latent factors to observed features in each data view, and (iii) a perturbation-to-factor coefficient matrix that quantifies how each perturbation modulates each latent program. A key design principle of PACMON is the incorporation of prior biological knowledge directly into the model structure. Specifically, pathway annotations from curated databases such as Hallmark, Reactome, and Gene Ontology are encoded as a structured sparsity prior on the loading matrices (Sec. 4.3) [Liberzon et al., 2015, Milacic et al., 2024, Ashburner et al., 2000]. This prior encourages each latent factor to align with a specific biological pathway by strongly shrinking loadings for genes outside the annotated gene set, while allowing genes within the pathway to contribute freely. Crucially, the regularised horseshoe prior permits genes outside the annotation to acquire non-zero loadings when strongly supported by the data, enabling data-driven refinement of pathway definitions beyond the initial prior knowledge [Qoku and Buettner, 2023]. The latent factors are modelled as functions of the experimental perturbations through a linear regression module (Sec. 4.2), such that each entry captures the signed effect of a given perturbation on each pathway program. This parametrisation directly links perturbations to interpretable biological programs rather than to individual genes, yielding a compact, low-dimensional summary of perturbation effects that facilitates downstream analyses including perturbation clustering, dose–response characterisation, and cross-condition comparison. To ensure biological interpretability, PACMON uses a nonnegative configuration for factor scores and loadings (Sec. 4.4), so that latent factors can be interpreted as additive biological programs with nonnegative feature contributions.

**Figure 1:**
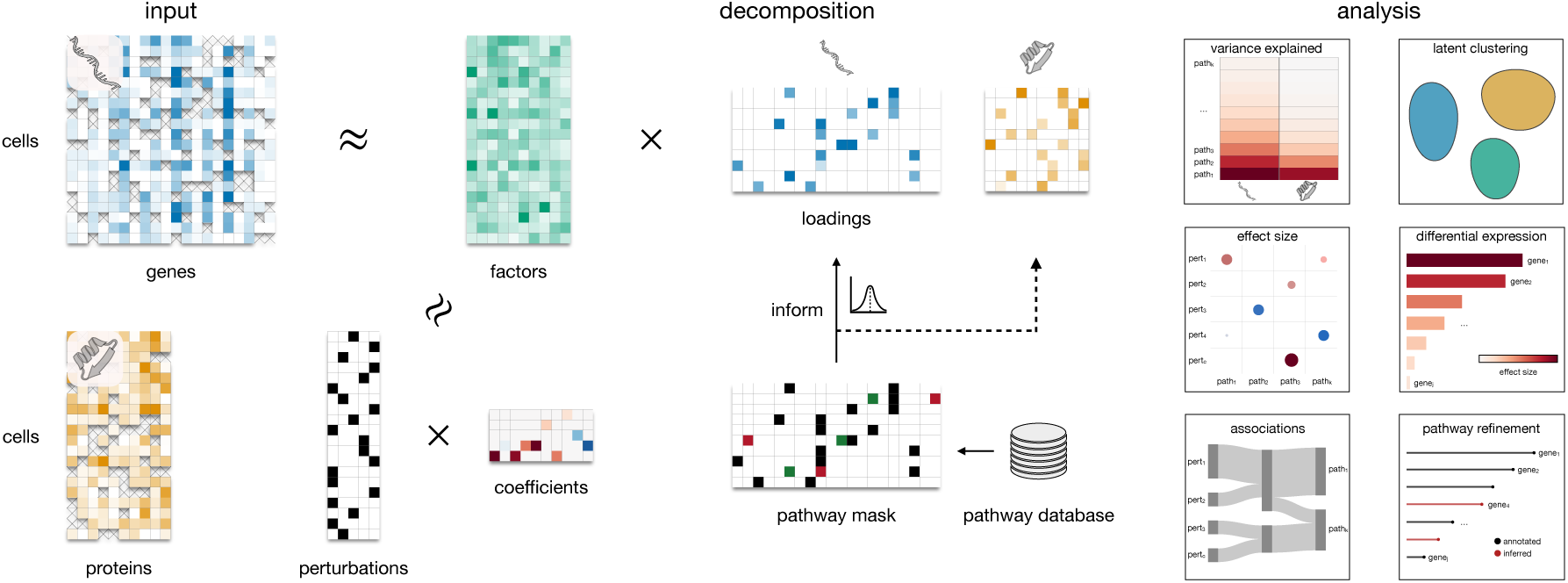
Overview of PACMON. Multimodal measurements, here RNA, surface proteins and perturbation identities, are decomposed into shared latent factors, modality-specific loadings and perturbation-to-factor coefficients under pathway-informed structured sparsity. This representation enables downstream analyses including variance decomposition, latent clustering, perturbation–pathway association analysis, differential expression and pathway refinement.

Statistical significance of inferred effects—including perturbation–pathway associations, perturbation–gene effects, and other derived quantities—is assessed through a unified posterior-significance framework based on the variational local false sign rate (vLFSR) combined with a region-of-practical-equivalence (ROPE) criterion (Sec. 4.6). The resulting representation enables a suite of downstream analyses (Fig. 1, right): variance decomposition across modalities to identify dominant sources of biological variation; latent clustering to reveal perturbation groupings in pathway space; perturbation–pathway association analysis to map how individual perturbations modulate specific biological programs; differential expression analysis by propagating perturbation effects through the loadings to obtain gene- and protein-level effect sizes (Sec. 4.7); and pathway refinement, whereby genes outside the original annotation that receive high data-driven loadings extend and contextualise curated pathway definitions.

### 2.2 PACMON outperforms specialised tools in terms of accuracy and scalability in a simulation study

We first evaluated PACMON on synthetic single-view perturbation data with known ground-truth latent factors, perturbation-factor coefficients, and downstream feature-level effects. To enable direct comparison with GSFA [Zhou et al., 2023], which operates on transcriptomic data only, we generated datasets with a single modality consisting of 10,000 samples, 5,000 features, 20 latent factors and 20 perturbation conditions. All methods were evaluated on the same simulated datasets across five random seeds. In order to more closely resemble real-world domain knowledge, we informed PACMON with noisy pathway information a priori. Specifically, 50% of the true positive features within each latent factor were swapped with randomly selected true negatives, thereby introducing both false positives and false negatives into the prior annotations. In addition, we included an uninformed baseline, denoted PACMON*_u_*, which simply places an uninformative Horseshoe prior on the factor loadings. PACMON accurately recovered both the latent pathway structure and the effects of perturbations on these latent programs. As shown in Fig. 2A, PACMON achieved near-perfect pathway recovery (AUPR = 0.99 ± 0.00) and the strongest recovery of perturbation effects (AUPR = 0.94 ± 0.04), while the uninformed PACMON*_u_*model also performed strongly (pathway recovery AUPR = 0.94 ± 0.04; perturbation-effect recovery AUPR = 0.88 ± 0.08). Both variants substantially outperformed GSFA, which achieved pathway and perturbation-effect AUPRs of 0.65 ± 0.05 and 0.60 ± 0.11, respectively. These results indicate that PACMON successfully recovers the modular perturbation structure of the data and benefits further from pathway-informed priors, even when those priors are substantially corrupted.

**Figure 2:**
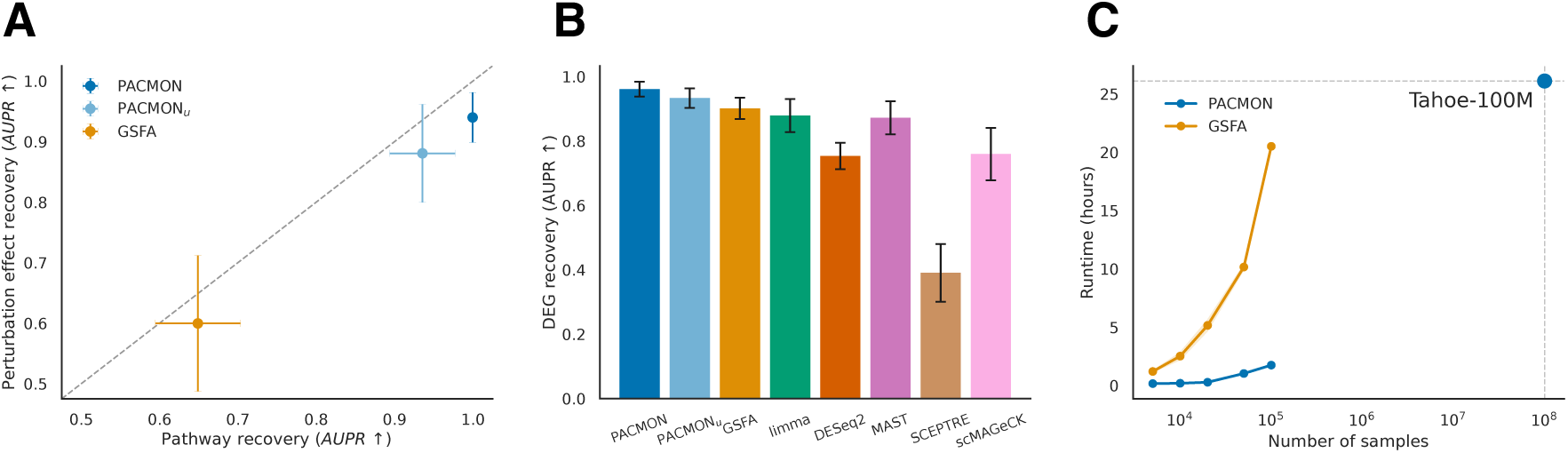
PACMON accurately recovers pathway and differential-expression signals while scaling to large perturbation datasets. **(A)** Recovery of underlying pathways and their perturbation-associated structure on synthetic data, measured by area under the precision–recall curve (AUPR), comparing PACMON with PACMON*_u_* (uninformed) and GSFA. PACMON achieves the highest recovery of ground-truth pathways, as well as their corresponding perturbation associations. **(B)** Recovery of differentially expressed genes on synthetic data, again measured by AUPR, showing strong agreement between PACMON and ground-truth perturbation-induced feature responses relative to alternative approaches. **(C)** Runtime as a function of dataset size, highlighting the scalability of PACMON relative to GSFA and enabling application to atlas-scale perturbation datasets such as Tahoe-100M.

We further validated the ability of PACMON to recover known effects of individual perturbations on pathways on an independent transcriptomic dataset with experimentally confirmed perturbation effects (Sec. 5.1, [Lalli et al., 2020]).

We next assessed recovery of downstream gene-level perturbation effects by comparing PACMON with six established differential expression (DE) approaches spanning bulk RNA-seq methods (DESeq2 [Love et al., 2014], limma [Ritchie et al., 2015]), single-cell-specific DE tools (MAST [Finak et al., 2015]), specialized tools CRISPR screen analysis (scMAGeCK-LR [Yang et al., 2020], SCEPTRE [Barry et al., 2021]), and the latent factor model GSFA [Zhou et al., 2023]. All methods were applied to the same simulated datasets to identify genes significantly affected by perturbations.

Summarised in Fig. 2B, PACMON achieved the highest AUPR (0.96 ± 0.02), followed by the uninformed variant PACMON*_u_* (0.93 ± 0.03) and GSFA (0.90 ± 0.03), outperforming limma, DESeq2, MAST, SCEPTRE and scMAGeCK under this simulation setting. Thus, modelling perturbation effects through latent pathway programs not only improves recovery of pathway-level structure, but also yields highly accurate downstream feature-level perturbation effects. Finally, we assessed computational scalability by varying dataset size while keeping the remaining simulation settings fixed. PACMON scaled substantially better than GSFA across all sample sizes tested. On a single Quadro RTX 5000 (16 GB), PACMON required on average 1.74 hours for 100,000 samples, whereas GSFA required 20.50 hours (Fig. 2C). This improved computational efficiency is consistent with the stochastic variational inference framework underlying PACMON, which is substantially more scalable than the Gibbs sampling inference used by GSFA [Zhou et al., 2023]. These results highlight the practical advantage of variational inference for large perturbation screens and enable the application of PACMON to atlas-scale single-cell data.

### 2.3 PACMON identifies pathway programs from multimodal perturbation data and reveals perturbation-pathway associations

We next applied PACMON to a multimodal CRISPR screen of patient-derived melanoma cells designed to identify tumour-intrinsic mechanisms of immune evasion [Frangieh et al., 2021]. This dataset contains measurements of both transcriptome and surface proteins from more than 150,000 single cells profiled using Perturb-CITE-seq across three experimental conditions: control, T cell co-culture, and IFN*γ* treatment. Cells carry single CRISPR perturbations targeting 241 genes or non-targeting control guides. To encourage interpretable latent structure, we informed PACMON with annotations from the Hallmark and Reactome databases. Specifically, PACMON ingested 26 Hallmark gene sets and 50 Reactome pathways. In addition to the pathway-informed latent factors, we included one additional dense latent factor, included by default in all PACMON analyses, that was excluded from the perturbation design, effectively acting as an intercept-like component that captures global variation not attributable to specific perturbations. We first examined the latent components explaining the dominant sources of variation in the data. Among the strongest gene programs we identified Interferon Signaling (R), Interferon Gamma Response (H), G2M Checkpoint (H), Cell Cycle Checkpoints (R), and E2F Targets (H) (Supplementary Fig. S1, S2). A UMAP representation of the joint latent space revealed clear separation of the three experimental conditions (Fig. 3A). IFN*γ*-stimulated cells were characterised by strong activation of the Interferon Gamma Response (H) program, whereas T cell co-culture induced increased activity of the broader Interferon Signaling (R) pathway [Frangieh et al., 2021] (Fig. 3C, S3). In addition to capturing condition-driven structure, PACMON also resolved intrinsic cell-state variation. Latent factors corresponding to cell-cycle programs displayed clear alignment with independently inferred cell-cycle phases (Supplementary Fig. S4, S5). In particular, the G2M Checkpoint (H) pathway showed strong activation in cells assigned to the G2/M phase, while the Reactome Cell Cycle Checkpoints pathway exhibited increased activity in S-phase cells (Supplementary Fig. S6). These results indicate that PACMON recovers biologically meaningful pathways that reflect both experimental perturbations and endogenous cell-state structure. The cell-cycle factors were further supported by coherent loading profiles: G2M Checkpoint (H) recovered canonical mitotic regulators and additionally highlighted *CCNB1* as a refined marker, whereas Cell Cycle Checkpoints (R) captured core DNA replication and S-phase genes (Supplementary Fig. S7). We next examined the composition of the inferred interferon-associated pathway factors to assess whether they captured known multimodal biology. The Interferon Gamma Response (H) factor exhibited strong RNA loadings for canonical interferon-stimulated genes including *IDO1*, *IRF1*, and *STAT1*, together with guanylate-binding proteins (*GBP1*, *GBP2*, *GBP4*) that are well-established interferon-inducible effectors [Tretina et al., 2019]. In the protein modality, this factor was dominated by PD-L1 (encoded by *CD274*) and also loaded on antigen-presentation and immune-interaction markers (e.g. HLA-A, CD47, CD58), consistent with IFN*γ*-driven checkpoint and visibility phenotypes [Moon et al., 2017] (Fig. 3E). Complementary to this, the broader Interferon Signaling (R) factor captured an antigen-presentation enriched transcriptional program, with strong RNA loadings on MHC class II components (*CD74*, *HLA-DRA*, *HLA-DRB1*, *HLA-DPA1/DPB1*, *HLA-DMA*), consistent with the IFN*γ*CIITA axis that induces MHC class II genes [Steimle et al., 1994, Léon Machado and Steimle, 2021] (Fig. 3F). These multimodal loading patterns indicate that PACMON recovers coherent interferon-driven programs spanning transcriptional interferon effectors, antigen presentation machinery, and surface immune checkpoint markers. Finally, we examined associations between perturbations and pathways inferred by PACMON to determine how genetic perturbations modulate pathway activity (Fig. 3D). Disruption of the IFN*γ*-JAK/STAT signaling cascade (*IFNGR1*, *IFNGR2*, *JAK1*, *JAK2*, *STAT1*) resulted in strong suppression of interferon-responsive programs, consistent with canonical resistance mechanisms [Frangieh et al., 2021]. PACMON captured pronounced negative associations with interferon-related pathways, reflecting a selective loss of interferon-driven latent activity rather than a global transcriptional effect. In line with this pattern, knockout of *CD274* (PD-L1) reduced activity of interferon-associated programs while modestly increasing scores for broader cytokine signaling pathways, suggesting a more nuanced inflammatory state. In contrast, perturbations affecting proliferative regulators such as *CCND1* primarily influenced pathways associated with cell-cycle progression, including Cell Cycle Checkpoints (R) and E2F Targets (H), while leaving interferon-linked programs largely unaffected. These results demonstrate that PACMON disentangles distinct biological axes and assigns perturbation effects to specific pathway programs rather than distributing them across unrelated transcriptional processes. Complete perturbation–pathway, perturbation–gene and perturbation–protein effect summaries are provided in Supplementary Figs. S8–S9 and Supplementary Tables S1–S3; a detailed analysis of STAT1 knockout effects at the gene and protein level is shown in Supplementary Figs. S11–S12.

**Figure 3:**
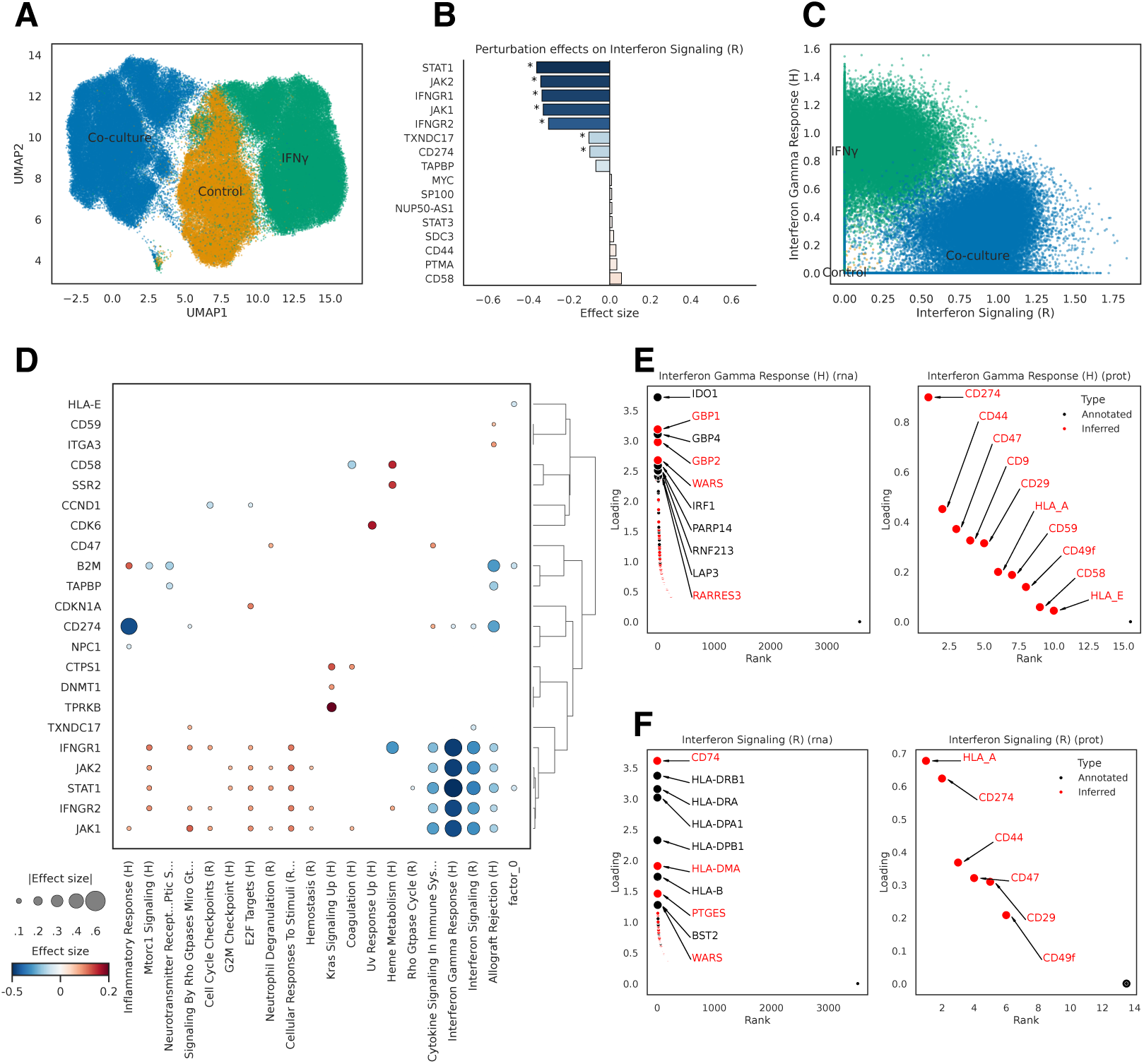
PACMON identifies interpretable pathway programs from multimodal perturbation data. **(A)** UMAP of the inferred joint latent space for the Perturb-CITE-seq melanoma dataset, showing separation of control, T cell co-culture and IFN*γ*-treated cells. **(B)** Perturbation effects on the Interferon Signaling (R) factor. Disruption of the IFN*γ*–JAK/STAT cascade strongly suppresses interferon-associated pathway activity, consistent with canonical immune-evasion mechanisms. **(C)** Joint distribution of Interferon Signaling (R) and Interferon Gamma Response (H) factor activities across cells, illustrating that IFN*γ* treatment is primarily associated with the Hallmark interferon-gamma program, i.e. Interferon Gamma Response (H), whereas co-culture shifts cells along a broader interferon-signaling axis. **(D)** Dotplot summarising perturbation–pathway associations across targeted genetic perturbations. Dot colour denotes effect direction and magnitude, and dot size denotes absolute effect size. The inferred landscape highlights pathway-specific responses across immune and proliferative programs. **(E)** Top RNA and surface-protein loadings for the Interferon Gamma Response (H) factor. PACMON recovers canonical interferon-stimulated genes together with immune-relevant surface proteins, including PD-L1 (*CD274*), indicating coordinated multimodal interferon responses. **(F)** Top RNA and surface-protein loadings for the broader Interferon Signaling (R) factor, showing enrichment of antigen-presentation machinery, including MHC class II components at the transcript level and immune-interaction markers at the protein level.

### 2.4 PACMON elucidates drug-response pathway landscapes in the Tahoe100M perturbation atlas

We applied PACMON to the Tahoe-100M single-cell perturbation atlas, which comprises approximately 100 million cells profiled across hundreds of chemical perturbations assayed at up to three concentrations (0.05, 0.5 and 5.0 µM) [Zhang et al., 2025]. The dataset contains 1,135 drug–dose combinations together with DMSO-treated vehicle controls—roughly three orders of magnitude larger than perturbation screens previously analysed with latent factor models and beyond the practical reach of existing perturbation-aware approaches such as GSFA (Fig. 2C). After training on a single GPU, PACMON produced a pathway-centric representation of drug-induced transcriptional responses. The inferred coefficient matrix encoding associations between perturbations and pathways exhibited a sparse structure (Supplementary Fig. S14), with most drug-dose perturbations showing supported effects on only a small subset of pathways. We summarised each drug by averaging pathway coefficients across the three assayed concentrations (0.05, 0.5 and 5.0 *µ*M) and visualised the most pronounced drug-pathway associations (two-sided tail quantile *q* = 0.005), yielding a compact subset of 36 drugs in Fig. 4A. The complete perturbation-pathway coefficient matrix is provided in Supplementary Table S4. To quantify alignment of inferred factors with their Hallmark annotations, we assessed enrichment of factor loadings using principal component gene set enrichment (PCGSE) [Frost et al., 2015]. All Hallmark-informed factors shown in Fig. 4 were significantly enriched for their corresponding gene programs after multiple-testing correction. The global drug-pathway landscape revealed coherent and interpretable response patterns across Hallmark programs (Fig. 4A). We first examined pathway structure along the axis defined by Interferon Gamma Response (H) and mTORC1 Signaling (H). In this projection, JAK/STAT inhibitors such as Tofacitinib and Fedratinib occupied the interferon-suppressed region, whereas mTOR inhibitors such as Rapamycin and Temsirolimus clustered along the mTORC1-suppressed axis (Fig. 4B). Drug-level hulls further showed how individual compounds traversed this pathway space across the three assayed concentrations (Supplementary Fig. S16). These patterns are consistent with the expected roles of JAK/STAT signaling in mediating interferon-responsive transcription and of mTORC1 signaling in regulating anabolic growth programs [Boor et al., 2017, Hu et al., 2021, Talpaz and Kiladjian, 2021, Le Tourneau et al., 2008, Li et al., 2013]. We next focused on stress-associated responses using the pathway space defined by UV Response Up (H) and Apoptosis (H). At the mechanism of action level, protein synthesis, proteasome, HDAC and CDK inhibitors occupied distinct but partially overlapping regions of this stress-apoptosis landscape (Supplementary Fig. S15). At the individual drug level, hulls across doses highlighted substantial heterogeneity in dose-dependent responses among representative compounds (Supplementary Fig. S17). In particular, the translation inhibitors Harringtonine and Homoharringtonine strongly activated UV Response Up (H), with Harringtonine additionally inducing Apoptosis (H), consistent with translation blockade eliciting cellular stress and cell-death programs [Tang et al., 2006]. Proteasome inhibitors produced a similarly strong stress-associated signature: Bortezomib and Ixazomib showed large positive UV Response Up (H) effects, consistent with proteotoxic stress and unfolded-protein-response engagement under proteasome blockade [Obeng et al., 2006]. Apoptosis-linked transcriptional programs were further highlighted by strong positive Apoptosis (H) effects for the CDK inhibitor Dinaciclib and the HDAC inhibitor Panobinostat [Gregory et al., 2015, Jeon et al., 2013]. To illustrate dose-resolved effects for an individual compound, we examined Dinaciclib across different concentrations. Supplementary Fig. S18 summarises the corresponding pathway-level responses, and Supplementary Fig. S19 shows the strongest gene-level effects at the highest assayed concentration (5.0 *µ*M). Together, these results illustrate how PACMON links dose-dependent perturbation effects at the pathway level to interpretable downstream gene expression changes. Finally, PACMON recovered and refined canonical Hallmark signatures, enabling multi-scale interpretation of drug effects at both the pathway and gene levels (Fig. 4C-F).

**Figure 4:**
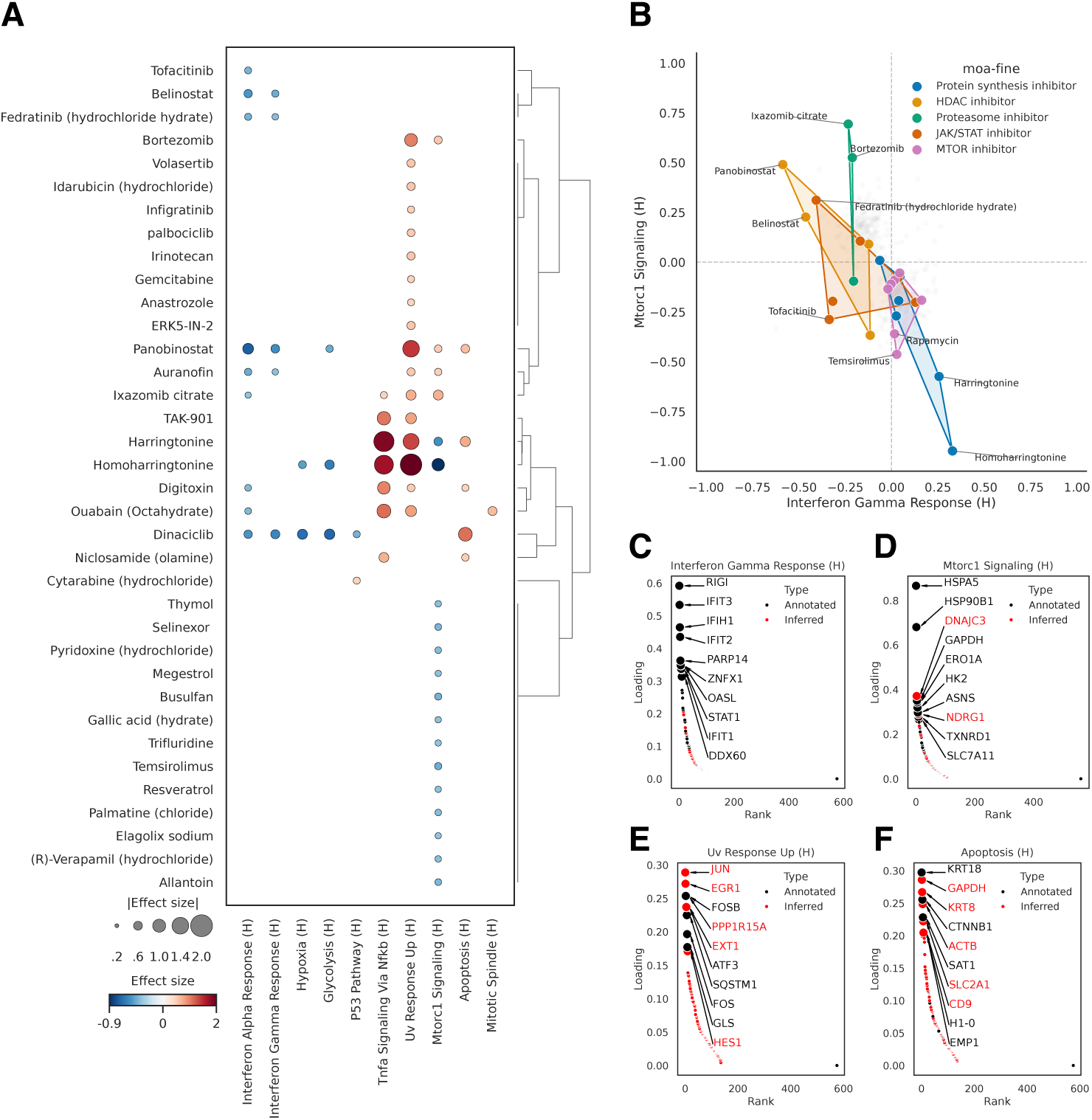
PACMON identifies interpretable drug-induced pathway programs in the Tahoe100M perturbation atlas. **(A)** Drug–pathway association landscape for the Tahoe-100M atlas. Each row corresponds to one drug and each column to a Hallmark pathway. Dot colour denotes effect direction and magnitude, and dot size denotes absolute effect size. For visualization, only extreme drug-pathway associations are shown after averaging coefficients across the three assayed concentrations, revealing coherent responses across interferon, growth, stress and apoptotic programs. **(B)** Projection of selected mechanisms of action in pathway space defined by Interferon Gamma Response (H) and mTORC1 Signaling (H). Each point corresponds to one drug, and convex hulls summarise the distribution of drugs within each mechanism of action class. JAK/STAT inhibitors occupy the interferon-suppressed region, whereas mTOR inhibitors cluster along the mTORC1-suppressed axis. **(C–F)** Top RNA loadings for representative pathway factors recovered from Tahoe-100M: Interferon Gamma Response (H) **(C)**, mTORC1 Signaling (H) **(D)**, UV Response Up (H) **(E)** and Apoptosis (H) **(F)**. Black points indicate genes present in the original Hallmark annotation, whereas red points denote genes inferred by PACMON outside the prior gene set. These loading profiles recover canonical pathway signatures while extending them with data-supported genes, highlighting the ability of PACMON to refine pathway definitions in atlas-scale perturbation data.

Together, these results illustrate how PACMON links dose-dependent perturbation effects at the pathway level to interpretable downstream gene-expression changes, enabling multi-scale interpretation from pathway programs to individual genes. Finally, PACMON recovered and refined canonical Hallmark signatures, extending prior gene-set definitions with data-supported genes at atlas scale (Fig. 4C–F; Supp. Fig. S13). Overall, these results demonstrate that PACMON scales to atlas-level perturbation data while maintaining pathway-level interpretability, enabling systematic mapping of drug-induced transcriptional programs across thousands of perturbations and ∼100 million cells.

## 3 Discussion and conclusion

We introduced PACMON, a Bayesian latent factor model that integrates multimodal molecular measurements with prior pathway knowledge to infer interpretable perturbation–pathway associations from large-scale perturbation screens. By combining pathway-informed structured sparsity with perturbation-aware latent modelling and scalable variational inference, PACMON addresses three key limitations of existing approaches: the lack of biological interpretability of inferred latent factors, the inability to jointly model perturbations and pathway programs, and poor computational scalability to modern atlas-scale datasets. PACMON is complementary to neural-network-based perturbation models that prioritise prediction of responses to unseen conditions [Lotfollahi et al., 2023a, Roohani et al., 2024, Zhu and Li, 2025]; PACMON instead provides an interpretable generative decomposition of observed perturbation effects with meaningful posterior uncertainty. On synthetic data, PACMON achieved near-perfect recovery of ground-truth pathway structure and perturbation effects, substantially outperforming GSFA in accuracy and runtime (Fig. 2). Notably, PACMON benefited from pathway-informed priors even when these were corrupted with 50% noise, and modelling perturbation effects through latent pathway programs improved recovery of gene-level differential expression relative to single-gene testing approaches. Validation on the LUHMES neurodevelopmental dataset further confirmed recovery of known perturbation–pathway relationships in a real biological context (Sec. 5.1).

PACMON is the first perturbation-aware latent factor analysis that allows to integrate multiple data modalities within a single generative model. Leveraging this integrative capability, we applied PACMON to a multimodal Perturb-CITE-seq melanoma dataset; it recovered biologically coherent interferon-signaling and cell-cycle programs spanning RNA and surface-protein modalities, and correctly identified disruption of the IFN*γ*–JAK/STAT cascade as the dominant driver of interferon-program suppression (Fig. 3). Application to the Tahoe-100M perturbation atlas across 100 million cells and over 1,000 drug–dose combinations yielded pharmacologically coherent drug-response landscapes across Hallmark pathway programs (Fig. 4). Existing perturbation-aware latent factor methods such as GSFA are limited to datasets roughly three orders of magnitude smaller, illustrating the practical necessity of the variational inference framework underlying PACMON.

Several limitations should be noted. First, PACMON currently assumes linear perturbation–factor relationships, which, while providing inherent interpretability, may not capture nonlinear effects; extending the model with nonlinear perturbation mappings could improve the modelling of combinatorial or nonlinear perturbation responses. Second, the Tahoe-100M analysis was restricted to 31 Hallmark programs for computational tractability. Finally, while the mean-field variational approximation provides substantial scalability advantages, exploring richer variational families could further improve model performance.

In conclusion, PACMON provides a unified, scalable and interpretable framework for mapping perturbation effects onto biological pathways from multimodal single-cell data. This enables a systematic characterisation of how genetic and chemical perturbations modulate coordinated biological programs from individual experiments to atlas-scale perturbation screens.

## 4 Methods

### 4.1 Factorisation model

PACMON is a Bayesian matrix factorisation method that decomposes high–dimensional molecular readouts from multiple modalities (data views) into a lower–dimensional latent space. For a single sample *i* ∈ {1*, . . . , N* }, e.g. single cell, let **y***_i_* ∈ ℝ*^D^* denote the *D*–dimensional molecular profile. In addition, assume the features are partitioned into *M* non–overlapping data views *G_m_* ⊆ {1, . . . , *D*} (e.g. different omics layers), such that *G_p_* ∩ *G_q_* = ∅ for *p* ≠ = *q*, |*G_m_*| = *D_m_*, and 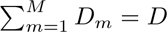. After reordering the features we write the data of view *m* as **Y**^(*m*)^ ∈ ℝ*^N^*^×*D*^*^m^*.

A core assumption of the Bayesian approach adopted for PACMON is that variation across all data modalities is jointly explained by a shared set of latent factors **z***_i_* ∈ ℝ*^K^* with *K* ≪ *D*. A linear mapping from the latent space to each modality is parameterised by the factor-loading matrix **W**^(*m*)^ ∈ ℝ*^Dm^*^×^*^K^* . The likelihood for cell *i* in view *m* can be written as

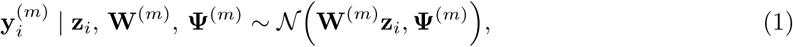

where **Ψ**^(*m*)^ = diag(***σ***^2(*m*)^) is a diagonal noise covariance with feature–specific residual variances 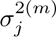. Collecting all latent factors into 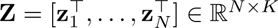, the induced matrix factorisation for each

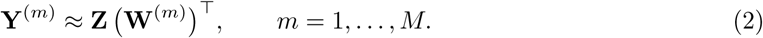

### 4.2 Perturbation–driven latent factors

Large–scale perturbation screens associate each cell with one or more experimental conditions, e.g., drug, dose, CRISPR perturbation, or their combinations. PACMON incorporates this structure by directly modelling how perturbations give rise to the latent factors.

Let **P** ∈ ℝ*^N^*^×*C*^ be a perturbation design matrix, where row **p***_i_* encodes the perturbation state of cell *i*. PACMON places a perturbation–informed prior on the latent factors:

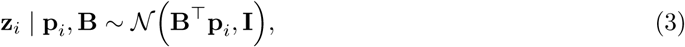

where **B** ∈ ℝ*^C^*^×*K*^ contains the perturbation–to–factor coefficients. The entry *B_c,k_* quantifies how strongly perturbation *c* activates (positive value) or represses (negative value) latent factor *k*. In addition to the perturbation-informed latent factors, PACMON includes one additional dense factor that is excluded from the perturbation design matrix, thereby acting as an intercept-like component that captures global variation not attributable to specific perturbations.

This parametrisation has two key consequences: (i) each latent factor corresponds to a perturbation–responsive gene program, and (ii) the matrix **B** yields a compact low–dimensional representation of perturbation effects, facilitating downstream tasks such as clustering of perturbations, dose–response characterisation, or cross–context comparison.

### 4.3 Sparsity and domain knowledge in the loadings

The loading matrices **W**^(*m*)^ define the mapping from latent factors to observed features and are therefore the main quantities used for biological interpretation. PACMON employs a regularised horseshoe prior to induce sparsity while simultaneously integrating prior knowledge in the form of noisy gene sets or pathways [Carvalho et al., 2009, Piironen and Vehtari, 2017, Qoku and Buettner, 2023].

For a generic loading element 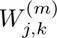 we specify

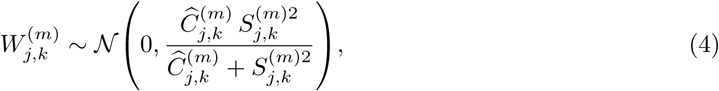

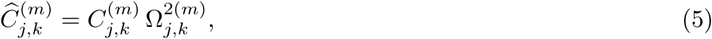

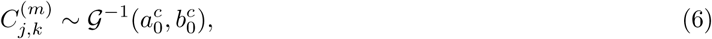

where 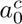 = 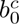 = 0.5 are fixed hyperparameters. The quantity

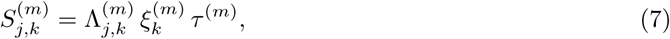

is the product of hierarchical shrinkage components,

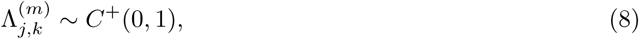

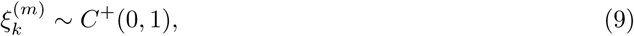

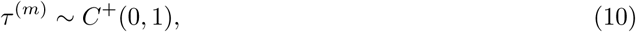

controlling element–wise, factor–wise, and view–wise shrinkage, respectively. The regularisation constants 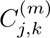 enforce a minimum amount of shrinkage even for large coefficients.

Prior knowledge is incorporated through a scaling matrix **Ω**^(*m*)^ ∈ [0, 1]^*D_m_×K*^. For an uninformed factor *k* we set 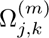 = 1 for all features *j*, recovering the standard regularised horseshoe prior. For an informed factor *k* we begin with a binary annotation (e.g. pathway membership or transcription factor regulon) and define

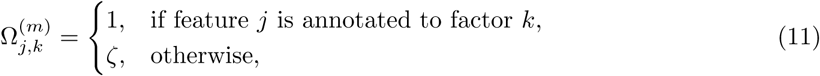

with a small relaxation constant *ζ* ≈ 0^+^ (we use *ζ* ≈ 0.01). Thus, features outside the annotated set receive a much stronger prior shrinkage, but may still acquire non-zero loadings when strongly supported by the data.

### 4.4 Non–negativity

For many single–cell applications it is desirable to interpret latent factors as additive gene programs whose activity and feature contributions are non–negative. PACMON adopts the non–negativity module introduced in MOFA-FLEX, which applies a rectified linear unit (ReLU) transformation to the latent variables, i.e., factor scores and factor loadings, during inference [Qoku et al., 2025].

Given unconstrained posterior samples of factor scores *Z_n,k_* and loadings 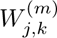, we define the transformed quantities

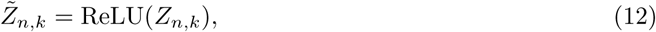

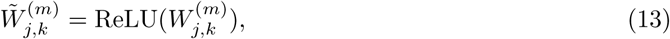

and replace **Z** and **W**^(*m*)^ with **Z̃** and **W̃**^(*m*)^ in the factorisation model of Equation (1).

This post–hoc transformation leaves the variational inference procedure and prior specification unchanged, but results in non–negative latent representations that can be interpreted as gene-activation programs and corresponding gene contributions. Importantly, PACMON allows non–negativity to be imposed independently on factor scores and loadings, providing flexibility for different biological applications.

### 4.5 Inference

Let 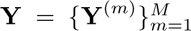 denote the observed multi–view data and let **P** ∈ ℝ*^N^*^×*C*^ be the observed perturbation design matrix. PACMON introduces latent factors **Z** ∈ ℝ*^N^*^×*K*^ that are generated from perturbations through the perturbation–to–factor coefficients **B** ∈ ℝ*^C^*^×*K*^ via

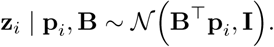

Each view *m* is generated through

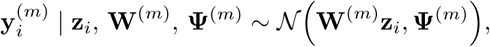

where **Ψ**^(*m*)^ = diag(***σ***^2(*m*)^). The weight matrices **W**^(*m*)^ follow the regularised horseshoe prior (Section 4.3).

The complete generative joint model factorises as

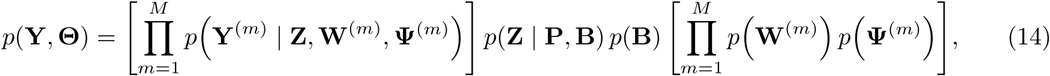

where the collection of all latent variables and parameters is

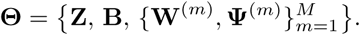

The full hierarchy of the regularised horseshoe prior on the loadings (Section 4.3) is included implicitly inside the term *p*(**W**^(*m*)^ | Λ^(*m*)^*, ξ*^(*m*)^*, τ* ^(*m*)^*, C*^(*m*)^, Ω^(*m*)^).

Since the posterior *p*(**Θ** | **Y**, **P**) is intractable, PACMON introduces a fully mean–field variational family

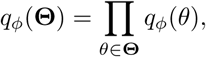

using Gaussian variational distributions for **Z**, **B** and each **W**^(*m*)^, and log–normal distributions for the positive scale parameters in the horseshoe hierarchy. The evidence lower bound (ELBO) is

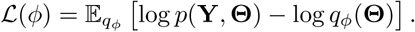

Optimisation is performed using stochastic variational inference. Monte–Carlo estimates of ∇*_ϕ_*𝓛 are computed using the reparameterization trick, ensuring low–variance gradient estimates. PACMON is implemented in Pyro, enabling automatic differentiation and scalable minibatch training.

### 4.6 Significance assessment of inferred effects

PACMON infers a range of effect sizes of interest, including perturbation-pathway associations, perturbation-gene effects, and other derived quantities obtained from linear transformations of latent variables and loadings. To assess the statistical significance and directionality of these inferred effects, we operate directly on their marginal variational posterior distributions. For each inferred effect, we use its variational posterior mean *µ* and standard deviation *σ*, and treat the effect as approximately Gaussian. Statistical evidence for a non-zero effect is quantified using the variational local false sign rate (vLFSR) [Stephens, 2017], defined as the posterior probability that the sign of the effect is incorrect,

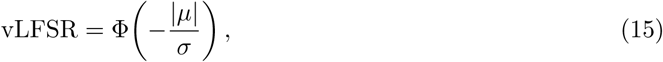

where Φ(·) denotes the cumulative distribution function of the standard normal distribution. Low vLFSR values indicate high confidence in the inferred sign of the effect. Since mean-field variational inference is known to underestimate posterior uncertainty, leading to inflated significant results, we apply a data-driven variance calibration procedure to improve uncertainty calibration without refitting the model [Wainwright and Jordan, 2008]. Posterior standard deviations are inflated by a global scaling factor estimated from the empirical distribution of |*µ*|*/σ*. Posterior standard deviations are inflated by a global scaling factor estimated from the central portion of the empirical distribution of |*µ*|*/σ*. By default, the central 60% of this distribution is retained and its median is matched to the median of a half-normal reference distribution. This calibration preserves relative effect sizes while reducing overconfidence in posterior estimates. To further guard against reporting effects that are statistically significant but negligible in magnitude, we apply a region-of-practical-equivalence (ROPE) criterion [Kruschke, 2013, 2018]. For an effect 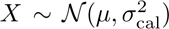, we compute the posterior probability that |*X*| exceeds a threshold *t*. By default, after variance calibration, the threshold is set to

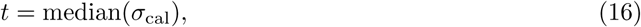

and effects are retained only if

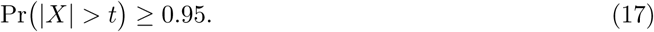

An effect is considered significant if it satisfies all of the following criteria: (i) vLFSR *<* 0.05, (ii) posterior mass outside the ROPE exceeds 0.95, and (iii) the sign of the effect is assigned according to the sign of the posterior mean. Significant effects are therefore reported in ternary form as positive, negative, or non-significant. This unified significance framework enables consistent statistical assessment across pathway-level perturbation effects, gene-level differential effects, and other downstream quantities derived from the PACMON model. All thresholds can in principle be adjusted, but the default settings were used throughout this study.

### 4.7 Derived perturbation effects on observed features

In addition to perturbation-factor coefficients, PACMON yields perturbation-feature associations for individual observed features by propagating perturbation effects through the view-specific loadings. For view *m*, let **B** ∈ ℝ*^C^*^×*K*^ denote the perturbation-factor coefficient matrix and **W**^(*m*)^ ∈ ℝ*^Dm^*^×^*^K^* the factor loading matrix. The induced perturbation-feature effect matrix is defined as

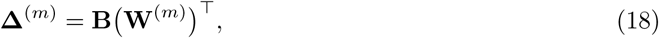

so that

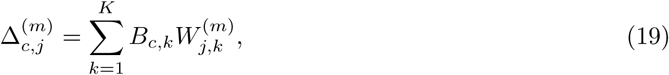

where 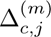 quantifies the signed effect of perturbation *c* on feature *j* in view *m*. Positive values indicate increased feature abundance under the perturbation, whereas negative values indicate decreased abundance. In the transcriptomic view, these quantities are interpreted as differential gene-expression effects, and in the protein view as differential protein abundance effects.

Posterior means for Δ^(*m*)^ are computed by multiplying posterior means of perturbation coefficients and factor loadings. Posterior variances are obtained under the mean-field approximation by summing the variances of the factor-wise products. Specifically, for each factor *k*, perturbation *c* and feature *j*, we use

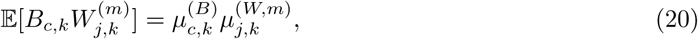

and

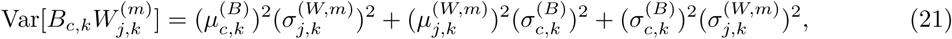

with the total variance of 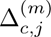 obtained by summing across factors *k*. Statistical significance for these derived perturbation-feature effects is assessed using the same posterior-significance framework described in Section 4.6.

### 4.8 Assessment of pathway alignment using PCGSE

To quantify agreement between inferred latent factors and pathway annotations, we adapted principal component gene set enrichment (PCGSE) [Frost et al., 2015] to the inferred factor loadings. For each selected view, we extracted the posterior mean factor loading matrix and restricted it to the features present in the tested feature sets. Loadings were rescaled by the maximum absolute loading within the view. Because PACMON was fit with the nonnegative configuration in the analyses presented here, we restricted the test to the positive part of the factor loading matrix. Posterior mean loadings of opposite sign, if present numerically, were set to zero, and the resulting positive loading magnitudes were used as feature-level statistics.

For each factor and feature set, we compared the mean loading magnitude of features inside the set against the mean loading magnitude of features outside the set using a two-sample *t*-statistic. Let *n*_in_ and *n*_out_ denote the number of features inside and outside the set, respectively, and let *w̅*_in_, *w̅*_out_, 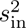 and 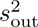 denote the corresponding sample means and variances of the loading magnitudes. We first formed the pooled variance estimate

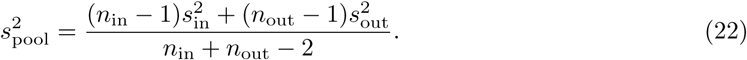

When correlation adjustment was enabled, we estimated the mean pairwise correlation among features inside the set from the observed data matrix of the corresponding view and defined a variance inflation factor

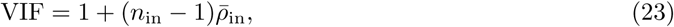

where 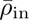 is the average within-set feature correlation. The resulting test statistic was

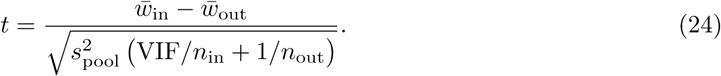

Two-sided *p*-values were obtained from the Student *t* distribution, and when one-sided tests were requested, effects with the opposite sign were set to zero and assigned *p* = 1.

Unless otherwise stated, only feature sets with at least 10 overlapping features were tested, correlation adjustment was enabled, and *p*-values were adjusted for multiple testing using the Benjamini-Hochberg procedure within each factor. Throughout the manuscript, pathway-informed factors with adjusted *p <* 0.05 were considered significantly aligned with their prior annotation. Informed factors that were not significant after correction were retained in the model but renamed as generic factors (for example, *factor 0*, *factor 1*) and interpreted as latent structure not confidently supported by the original pathway priors.

### 4.9 Data preprocessing and model setup

#### 4.9.1 Perturb-Seq Data

We first applied standard single-cell quality control procedures using Scanpy [Wolf et al., 2018]. Cells with fewer than 200 detected genes, mitochondrial transcript fractions exceeding 18%, or extreme library sizes outside the 1st-99th percentile of total UMI counts were removed. To enable downstream assessment of pathways associated with the cell cycle, we annotated cell-cycle states using curated S-phase and G2/M-phase gene sets following established protocols [Satija et al., 2015]. CRISPR guide annotations were parsed to determine the targeted gene for each cell. Non-targeting guides were collapsed into a single control category, and cells containing multiple distinct gene perturbations were excluded to retain only single-gene perturbations or non-targeting controls. To ensure robust estimation of perturbation effects, we further restricted the dataset to perturbations represented by at least 30 cells. Finally, we identified highly variable genes using a Seurat-style variance stabilisation procedure with variance estimation performed separately across experimental conditions and retained the top 6,000 genes for downstream modeling. All 20 surface proteins measured by the CITE-seq assay were retained.

#### 4.9.2 Tahoe-100M

We preprocessed each experimental plate independently following recommendations of the original study [Zhang et al., 2025]. We selected the top 10,000 highly variable genes per plate and intersected these sets across the 14 plates, resulting in a shared set of 4,148 genes. From this common feature space, we filtered the Hallmark gene set collection to retain programs with sufficient representation in the data (at least 20% of genes and at least 25 genes per program), yielding 31 gene programs. Restricting the expression matrix to genes belonging to these programs resulted in a final feature space of 1,014 genes used for model fitting. To prioritise pathway interpretability, we set a strong prior confidence (1 − *ζ* with *ζ* = 10^−8^) for the 31 Hallmark programs. Model fitting was performed via stochastic variational inference using minibatches of 10,000 cells, and we employed the scDataset framework for lazy on-disk batch loading [D’Ascenzo and Di Montesano, 2025] to reduce data-loading overhead. Training on a single NVIDIA H200 NVL GPU completed in approximately 25 hours.

#### 4.9.3 LUHMES data

The LUHMES dataset, which is publicly available on the Gene Expression Omnibus with the accession number GSE142078, consists of 8,843 cells and 33,694 genes, which were measured in three different runs. The Scanpy package [Wolf et al., 2018] was used to preprocess the data. Initially, cells with more than 10% mitochondrial RNA were removed. Furthermore, we filtered for cells with fewer than 20,000 total counts. We normalised to a target sum of 10,000 counts per cell and pseudo-log-transformed the counts. Finally, we selected the top 6,000 highly variable genes. During training, we corrected for total and mitochondrial counts, as well as for the different experimental runs, to account for batch effects. We used the publicly available Hallmark and GOBP gene sets from MSigDB [Liberzon et al., 2011, 2015, Ashburner et al., 2000] as prior annotations for model training. As we were particularly interested in pathways related to neuronal development, we selected GOBP pathways containing the substrings `NEUR, NERV, SYNAP`, and `AXON`. We filtered these pathways for the highly variable genes in the preprocessed dataset and set the minimum fraction of genes in a pathway that must be present in the dataset to 30%. The neuron-related pathways were then filtered further by setting a minimum count of 60 genes, in order to reduce the total number of prior pathways and to avoid many overlapping gene sets.

### 4.10 Aggregation and visualisation of drug-level pathway effects

For the Tahoe-100M analysis, perturbation effects were first inferred at the level of individual drugdose combinations. Drug-level summaries were then obtained by averaging pathway coefficients across the three assayed concentrations (0.05, 0.5 and 5.0 *µ*M). For visualisation of the global drug-pathway landscape, only the most extreme entries were retained using a two-sided tail quantile threshold of *q* = 0.005. Mechanism-of-action summaries were obtained by grouping drugs according to the annotated mechanism classes, whereas drug-level hull plots retained the individual dose-specific responses.

## Code availability

An open-source implementation of PACMON is available on GitHub at https://github.com/mlo-lab/ pacmon, where users can find detailed installation instructions as well as extensive tutorials.

## Acknowledgements

This work was supported by the Deutsche Forschungsgemeinschaft (DFG, German Research Foundation) – TRR 387/1 – 514894665 (T.S.).

## 5 Supplementary Results

### 5.1 PACMON correctly links *ASD* perturbations to biological processes

As an additional transcriptomic-only application, we validated PACMON on the Lund human mesencephalic (LUHMES) single-cell RNA-seq perturbation dataset [Lalli et al., 2020]. In this study, 14 genes associated with autism spectrum disorder (*ASD*) were perturbed in LUHMES neural progenitor cells to assess their effects on neuronal differentiation. The primary analysis revealed that perturbations of *CHD2*, *ASH1L*, *ARID1B*, and *DYRK1A* delay neuronal differentiation, whereas perturbations of *PTEN* and *CHD8* promote differentiation [Lalli et al., 2020]. To assess whether PACMON could recover these perturbation effects in an end-to-end manner, we ran PACMON with Hallmark and GOBP gene sets as prior pathway annotations. We first examined whether the inferred perturbation effects preserved the known structure among *ASD*-associated perturbations. To this end, we computed cosine similarity between perturbation-factor coefficient vectors for the factors explaining 98% of the variance. Supplementary Fig. S20 shows that *PTEN* and *CHD8* have highly similar perturbation profiles, whereas perturbations previously associated with delayed neuronal differentiation, namely *CHD2*, *ASH1L*, *ARID1B*, and *DYRK1A*, form a more distinct group. This organisation is consistent with the developmental phenotypes reported in the original study. We next examined perturbation-pathway associations in more detail. PACMON identified six pathways associated with at least one of the *ASD*-related perturbations (Supplementary Fig. S21). One example is a factor related to synapse organisation, which showed a negative association with *PTEN* perturbation. This is consistent with the established role of *PTEN* in synaptic regulation and plasticity [Sperow et al., 2012]. A second example factor is related to neuron development and showed positive associations with *PTEN* and *CHD8* perturbations and a negative association with *ASH1L* perturbation. These directions are biologically plausible: *PTEN* perturbation has been linked to increased proliferation and growth [Lalli et al., 2020, Chen et al., 2015], *CHD8* regulates multiple aspects of brain development [Sugathan et al., 2014], and *ASH1L* perturbation has been associated with altered neuronal morphogenesis [Cheon et al., 2022]. Inspection of the corresponding factor loadings provided further support for the biological relevance of these programs (Supplementary Fig. S22). For the neuron-development-associated factor, many of the top-ranked genes were part of the original GOBP annotation, but PACMON also highlighted additional genes outside the prior set. Among these, *NEFM* is a neurofilament protein implicated in neurite formation [Lariviere and Julien, 2004], and together with *NEFL* contributes to axonal cytoskeleton development [Yuan et al., 2017]. The factor also highlighted *MAP2* and *MAP1B*, which are important for microtubule organisation during neuronal development [Teng et al., 2001]. These results illustrate that PACMON not only recovers known biologically meaningful perturbation-pathway associations, but can also refine pathway definitions by assigning high weight to genes with relevant developmental functions that were not explicitly included in the original annotation.

## 6 Supplementary Figures

**Figure S1:**
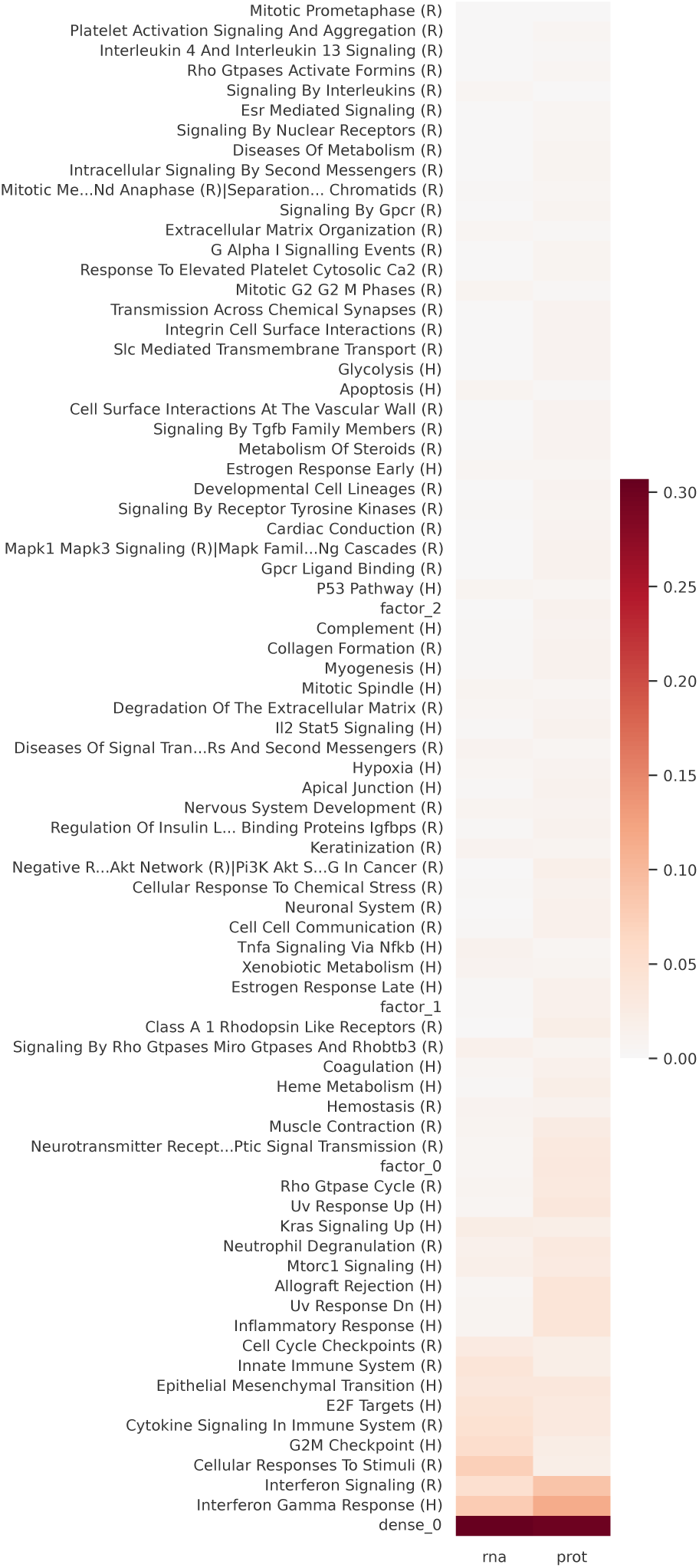
Variance explained by inferred factors across RNA and protein modalities in the PerturbCITE-seq melanoma dataset. Heat map showing the fraction of variance explained (*R*^2^) by each inferred latent factor in the RNA and protein views. Rows correspond to pathway-informed factors from the Hallmark and Reactome collections together with additional latent factors. Interferon-related and cell-cycle-associated programs account for substantial variance across both views, whereas other factors show stronger modality-specific contributions. The factor dense 0 denotes the uninformed latent factor that is excluded from the perturbation design and captures residual variation independent of specific perturbations. In contrast, factor_0, factor_1, and factor_2 correspond to previously pathway-informed factors that were not retained as significantly aligned with their prior annotations by PCGSE, indicating latent structure that is captured by the model but not confidently attributable to the original pathway priors.

**Figure S2:**
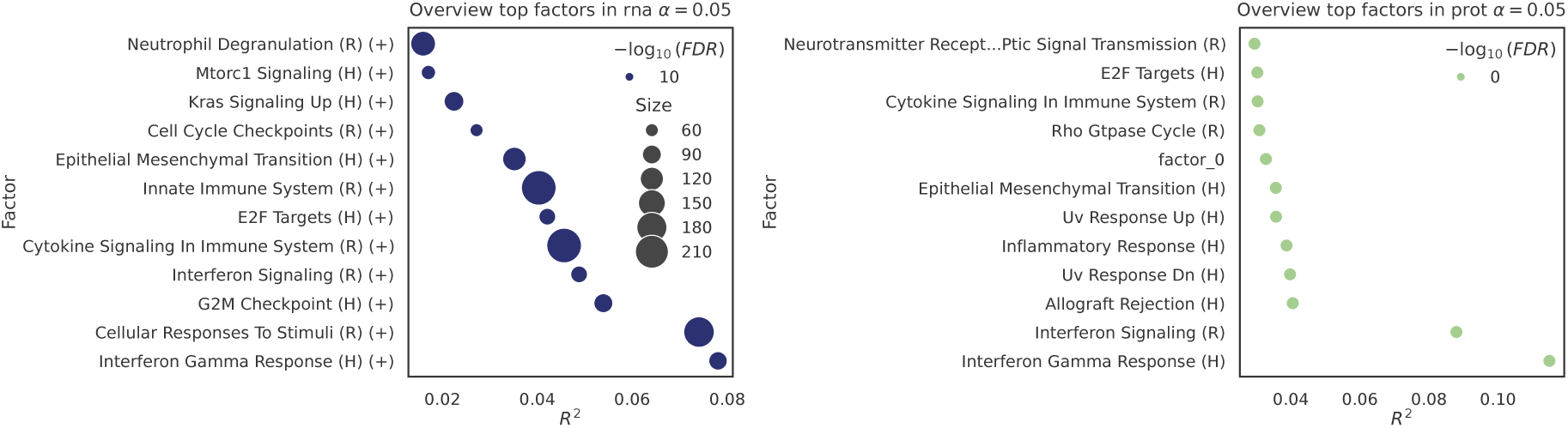
Overview of top latent factors in the multimodal Perturb-CITE-seq melanoma dataset. Bubble plots summarising the top factors identified by PACMON in the RNA (**left**) and protein (**right**) modalities at FDR < 0.05. The x-axis shows the fraction of variance explained (*R*^2^) by each factor, bubble size indicates factor size, and colour intensity corresponds to − log_10_(FDR). In the RNA view, dominant factors include interferon-related, cytokine-signaling, innate-immune and cellcycle programs, whereas the protein view is dominated by interferon-associated and inflammatory programs together with epithelial-mesenchymal transition and UV/stress-related signatures. The additional factor factor 0 denotes latent structure recovered by the model that was not significantly aligned with its original pathway annotation by PCGSE.

**Figure S3:**
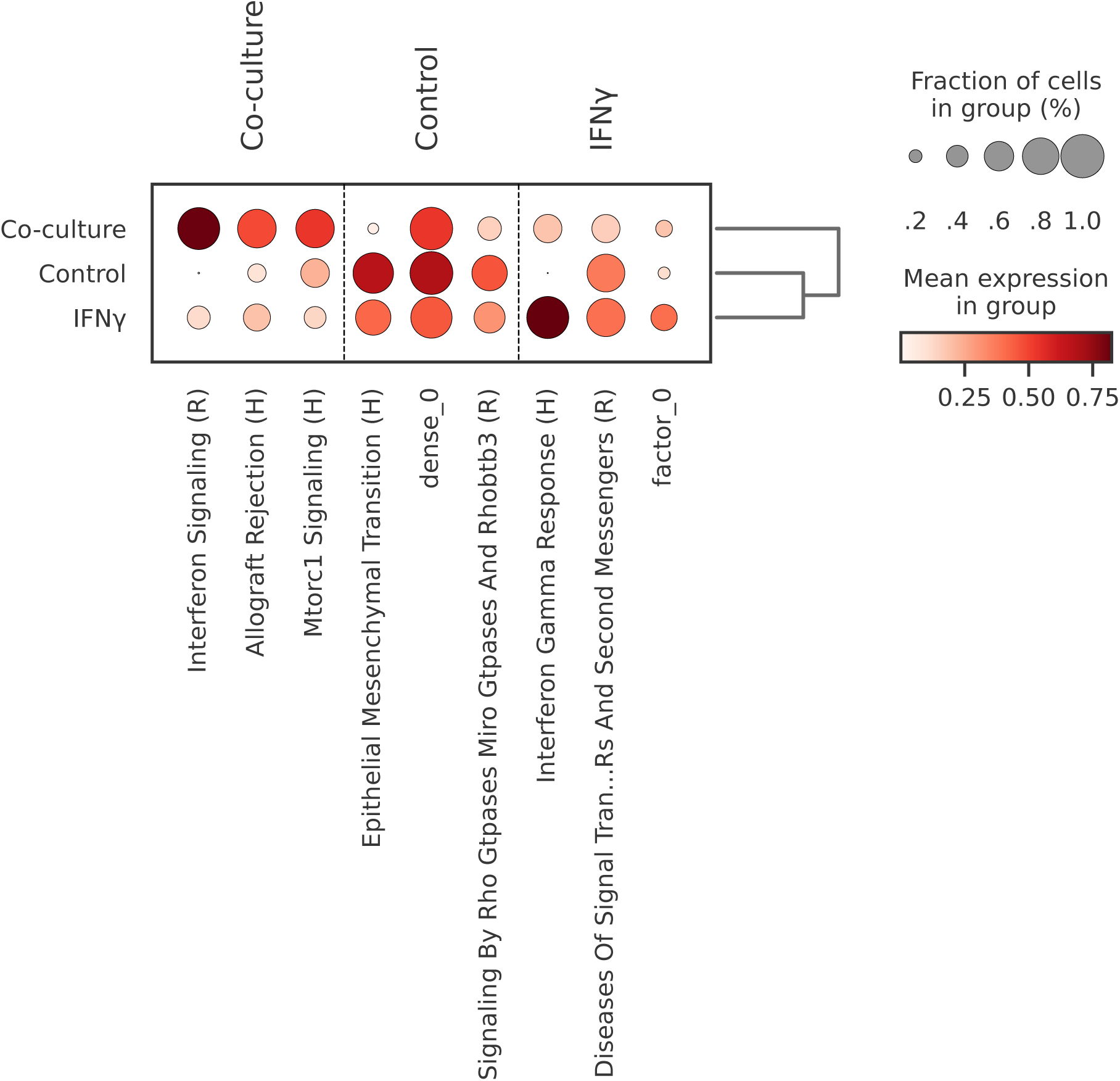
Differential pathway activity across experimental conditions in the Perturb-CITE-seq melanoma dataset. Dotplot summarising pathway activity differences between each condition and the remaining cells, assessed using Wilcoxon rank-sum testing. Dot size denotes the fraction of cells in the group, and colour indicates the mean pathway activity in that group. The co-culture condition is primarily associated with increased activity of Interferon Signaling (R), whereas IFN*γ*-treated cells are most strongly characterized by Interferon Gamma Response (H).

**Figure S4:**
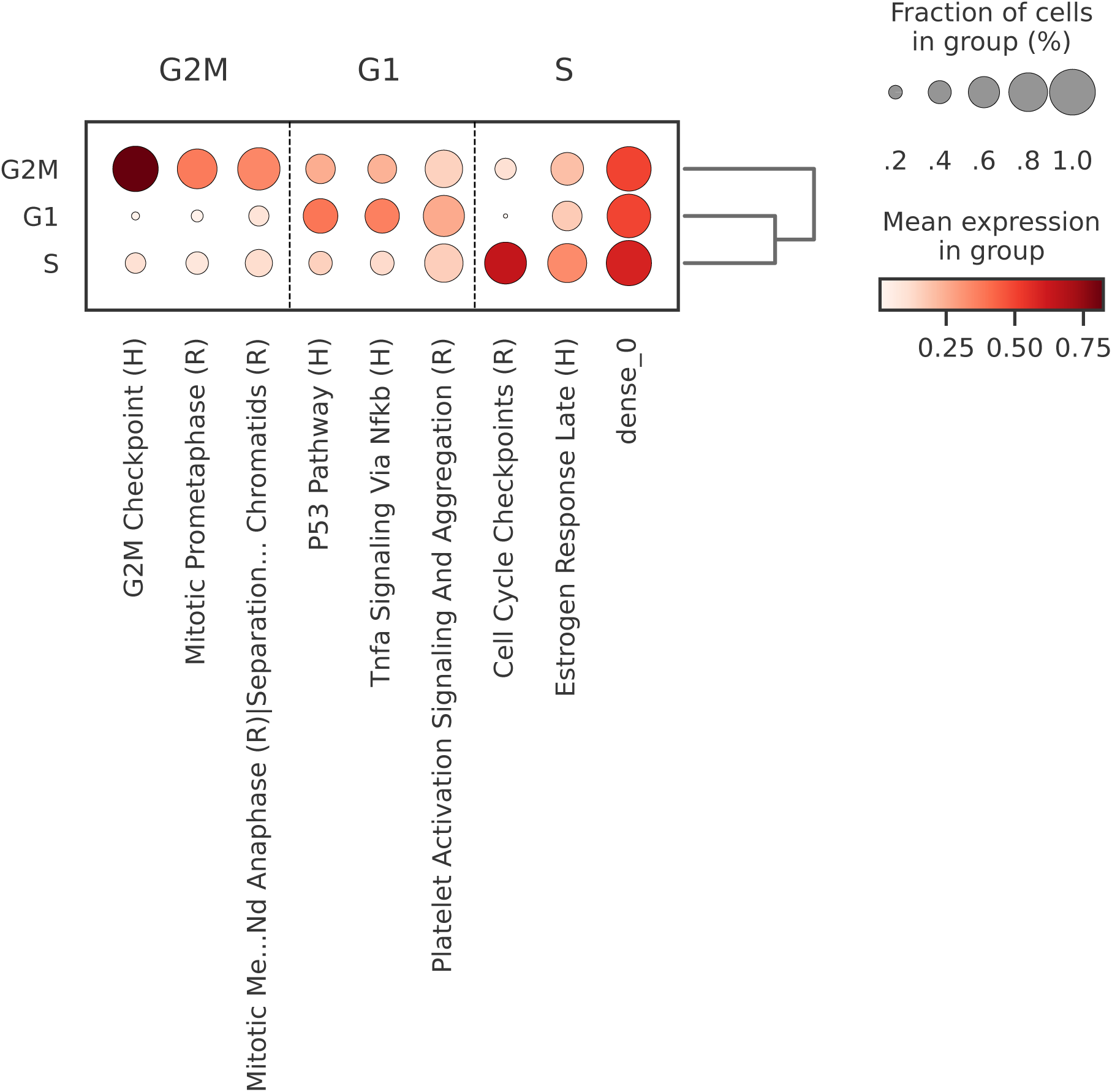
Differential pathway activity across cell-cycle phases in the Perturb-CITE-seq melanoma dataset. Dotplot summarising pathway activity differences between each cell-cycle phase and the remaining cells, assessed using Wilcoxon rank-sum testing. Dot size denotes the fraction of cells in the group, and colour indicates the mean pathway activity in that group. The G2/M phase is most strongly associated with G2M Checkpoint (H), whereas S-phase cells are characterized by increased activity of Cell Cycle Checkpoints (R). G1-phase cells additionally show elevated activity of P53 Pathway (H).

**Figure S5:**
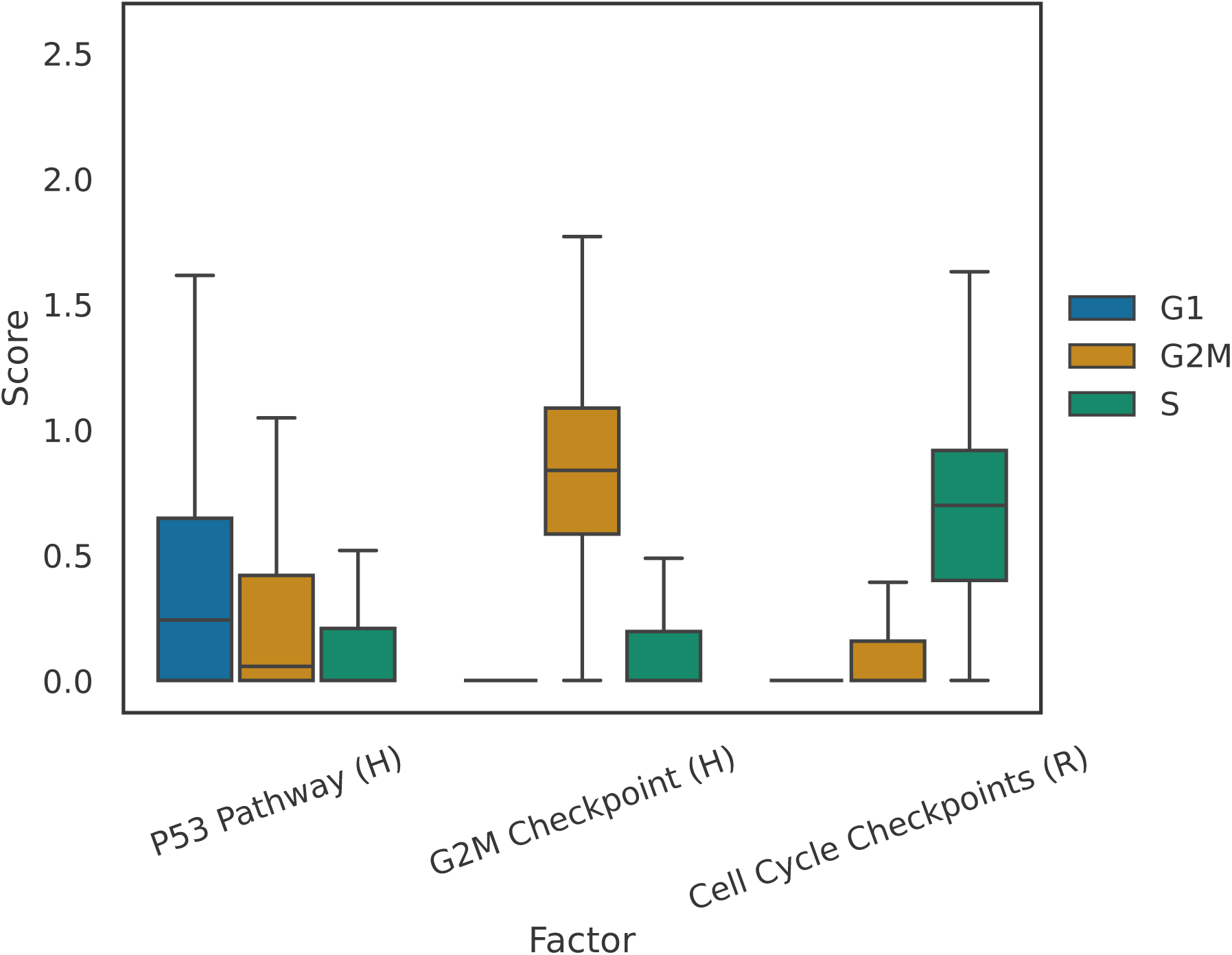
Cell-cycle-associated pathway scores across independently assigned cell-cycle phases in the Perturb-CITE-seq melanoma dataset. Boxplots showing the distribution of inferred pathway scores for P53 Pathway (H), G2M Checkpoint (H) and Cell Cycle Checkpoints (R) across cells assigned to G1, S and G2/M phases. Consistent with the differential testing results, G2M Checkpoint (H) is highest in G2/M-phase cells, Cell Cycle Checkpoints (R) is highest in S-phase cells, and P53 Pathway (H) shows elevated activity in G1-phase cells.

**Figure S6:**
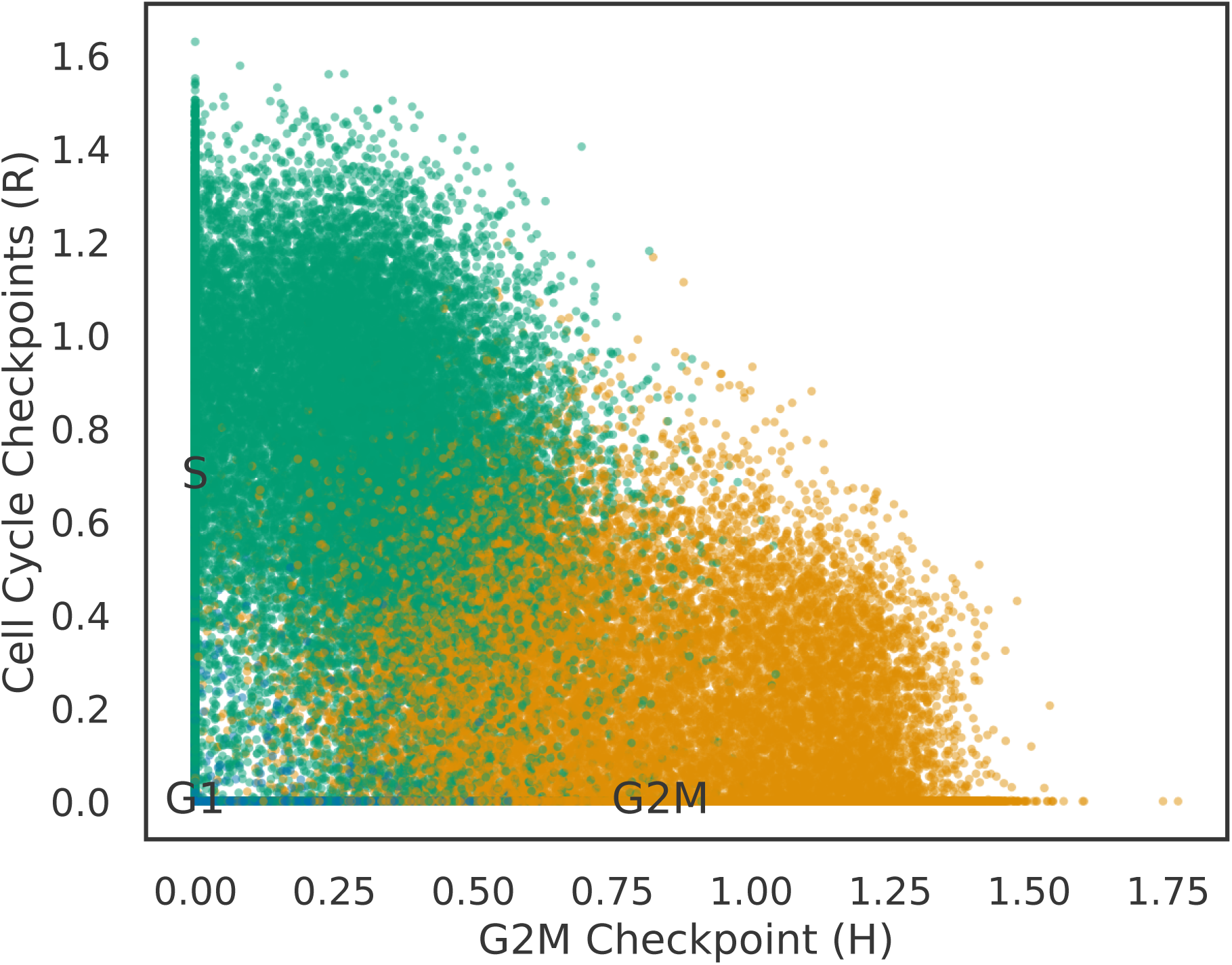
Joint distribution of G2/M and S-phase pathway activities in the Perturb-CITE-seq melanoma dataset. Scatterplot of factor scores for G2M Checkpoint (H) and Cell Cycle Checkpoints (R), coloured by independently assigned cell-cycle phase. Cells in G2/M preferentially occupy higher G2M Checkpoint (H) scores, whereas S-phase cells are enriched for high Cell Cycle Checkpoints (R) scores. G1-phase cells remain concentrated near the origin, consistent with low activity of both cellcycle programs outside actively cycling states.

**Figure S7:**
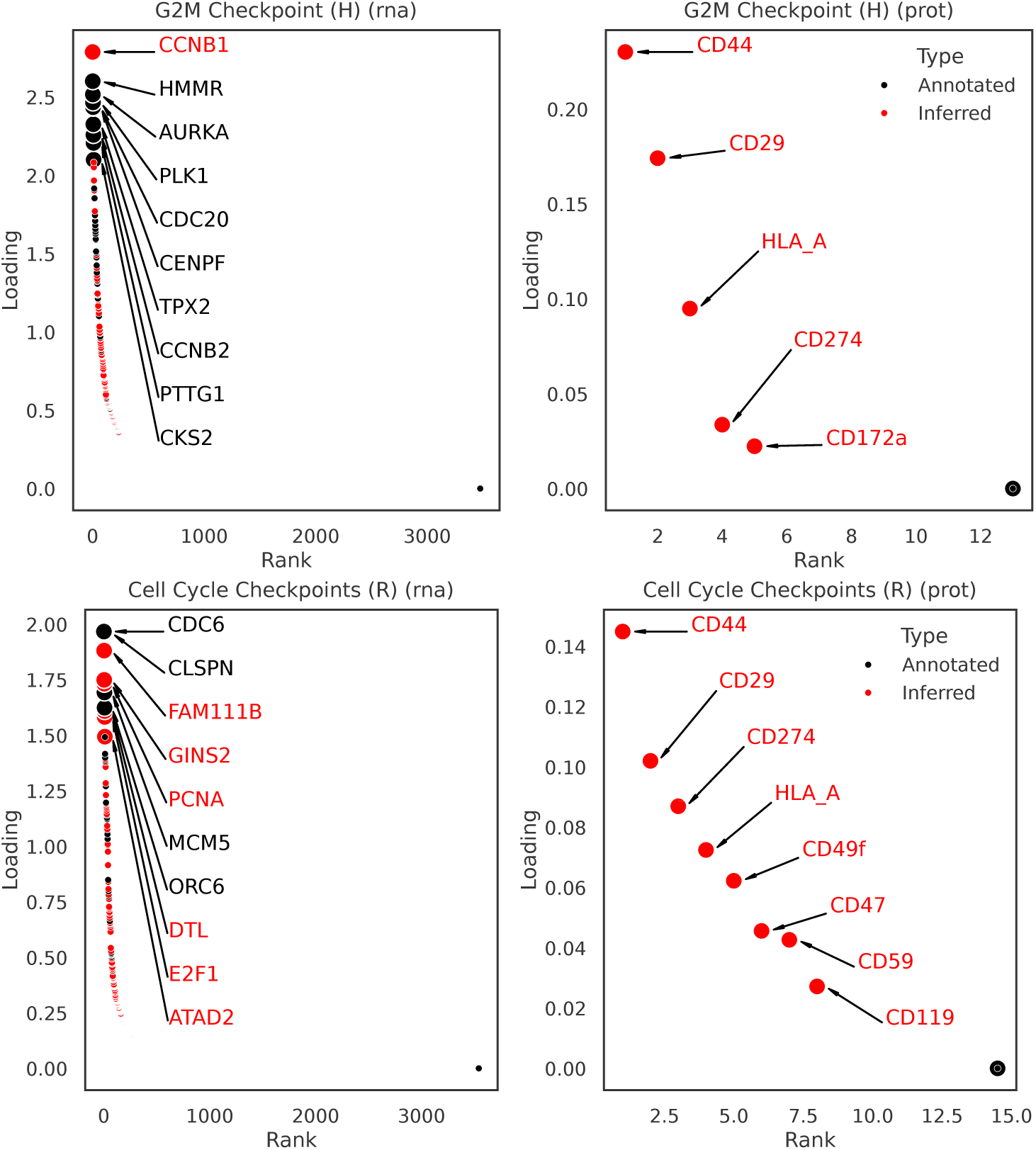
RNA and protein loadings for cell-cycle-associated pathway factors in the Perturb-CITEseq melanoma dataset. Top-ranked loadings for G2M Checkpoint (H) (**top**) and Cell Cycle Checkpoints (R) (**bottom**) are shown for the RNA (**left**) and protein (**right**) modalities. Black points denote genes present in the original pathway annotation, whereas red points indicate features inferred by PACMON outside the prior set. For G2M Checkpoint (H), the RNA loadings recover canonical mitotic regulators including *HMMR*, *AURKA*, *PLK1*, *CDC20*, *CENPF* and *TPX2*, while additionally assigning high weight to *CCNB1* despite its absence from the original annotation, indicating biologically meaningful pathway refinement. For Cell Cycle Checkpoints (R), the RNA loadings highlight core DNA replication and S-phase regulators such as *CDC6*, *CLSPN*, *MCM5* and *ORC6*, together with refined markers including *FAM111B*, *GINS2*, *PCNA*, *DTL*, *E2F1* and *ATAD2*. Corresponding protein loadings show that both factors are also reflected in the surface-protein modality, indicating that PACMON captures coherent cell-cycle-associated structure across RNA and protein readouts.

**Figure S8:**
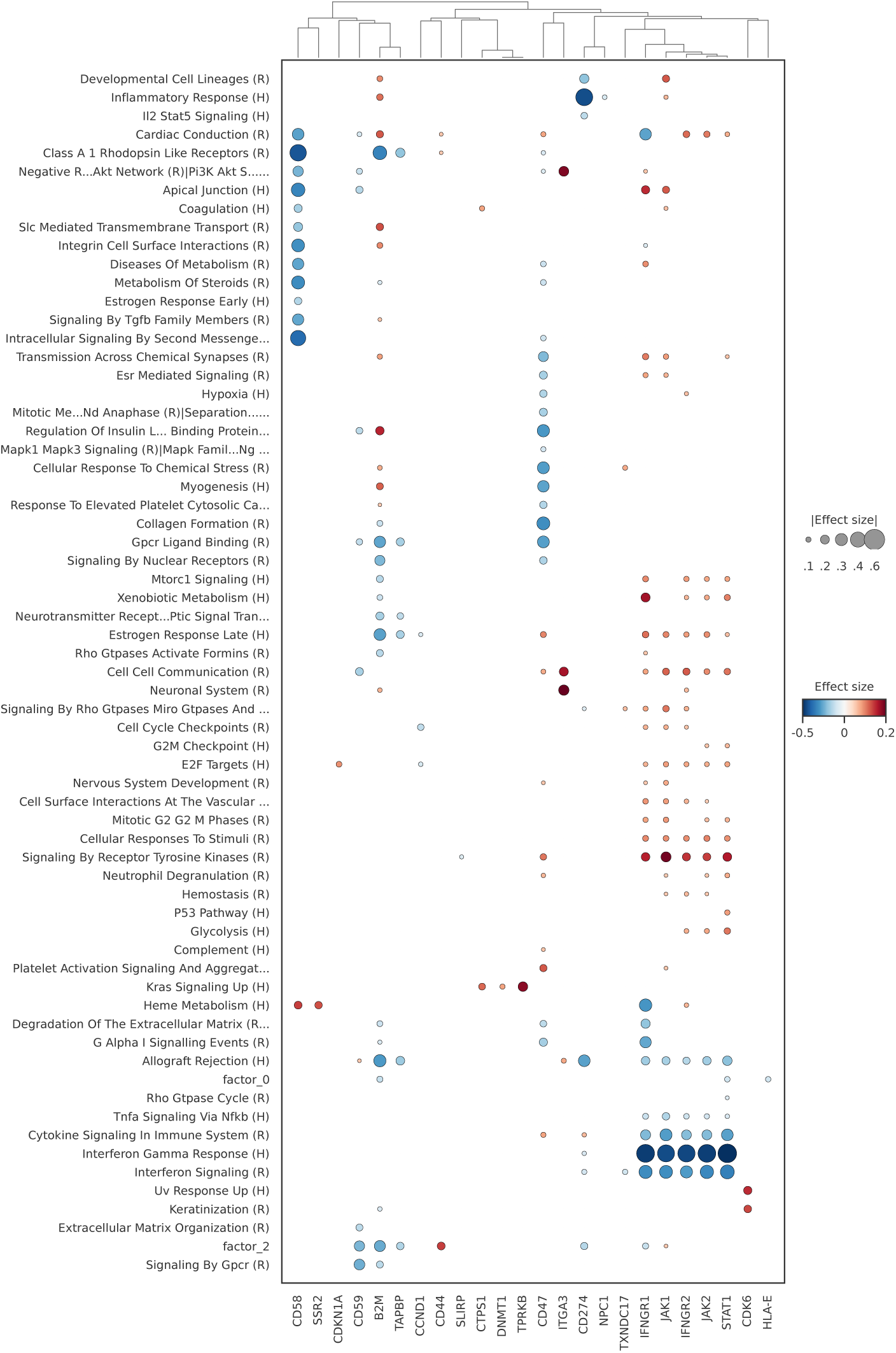
Global perturbation-pathway association landscape in the Perturb-CITE-seq melanoma dataset. Full dotplot of inferred perturbation-pathway associations, restricted to pathways with at least one significant perturbation effect and perturbations with at least one significant pathway association. Dot colour denotes effect direction and magnitude, and dot size denotes absolute effect size. This overview provides a global view of the pathway-level structure recovered by PACMON across the full set of significant perturbation responses.

**Figure S9:**
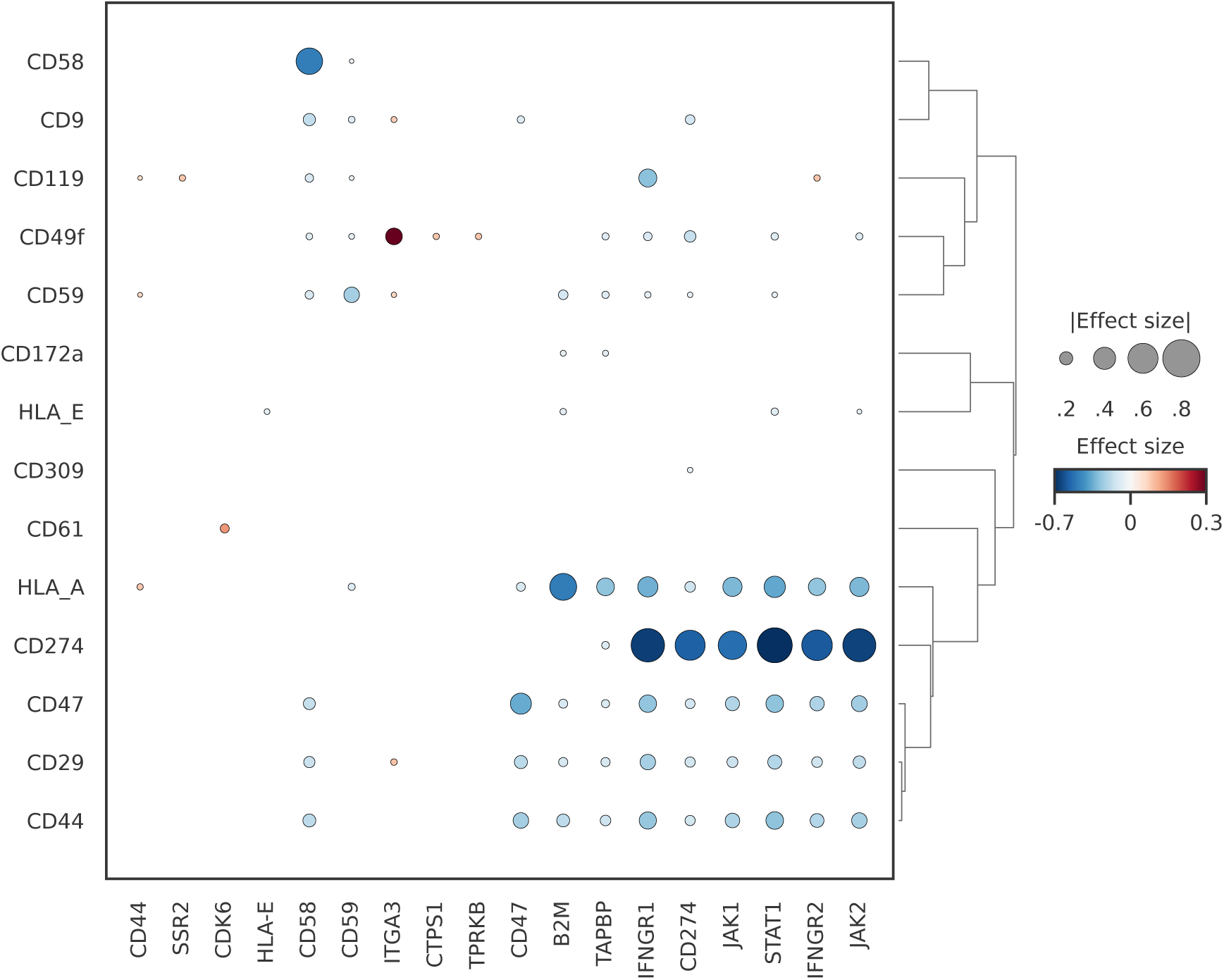
Perturbation-associated surface-protein effects that are differentially expressed in the Perturb-CITE-seq melanoma dataset. Dotplot summarising significant perturbation effects on surface-protein abundance. Rows correspond to measured proteins and columns to perturbations, with dot colour denoting effect direction and magnitude and dot size indicating absolute effect size. Consistent with the pathway-level analysis, perturbations targeting the IFN*γ*-JAK/STAT axis induce coordinated changes across multiple immune-relevant surface markers, including strong suppression of *CD274* (PD-L1), highlighting the ability of PACMON to recover interpretable perturbation effects directly in the protein modality.

**Figure S10:**
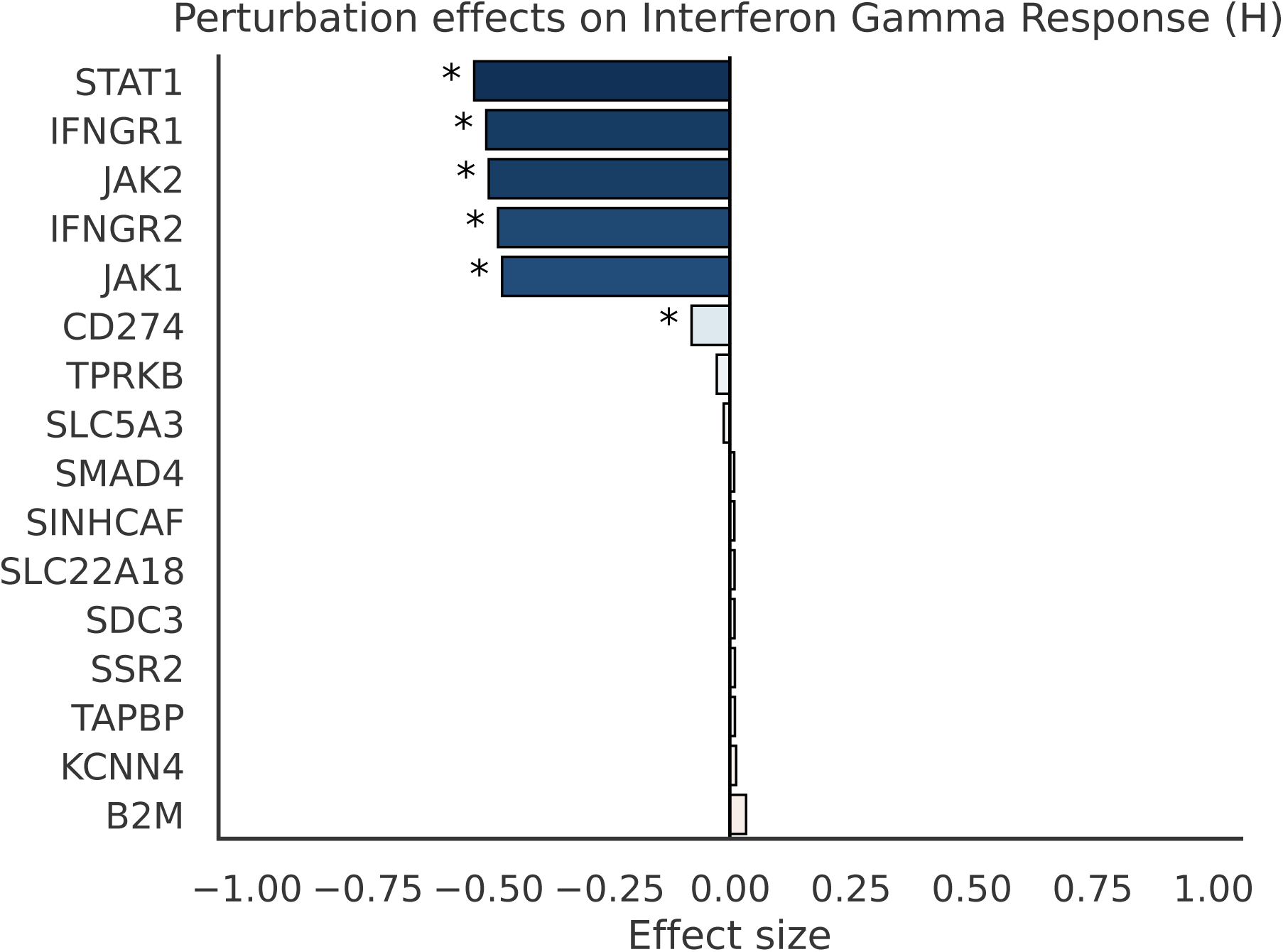
Perturbation effects on the Interferon Gamma Response (H) factor in the Perturb-CITEseq melanoma dataset. Barplot showing inferred perturbation effects on the Interferon Gamma Response (H) factor. Negative effect sizes indicate suppression of interferon-associated pathway activity, whereas positive values indicate increased activity. Consistent with the pathway-level dot plot, disruption of the IFN*γ*-JAK/STAT cascade, including *STAT1*, *IFNGR1*, *IFNGR2*, *JAK1* and *JAK2*, produces the strongest negative effects on this factor. Knockout of *CD274* (PD-L1) also reduces Interferon Gamma Response (H) activity, whereas the remaining perturbations show comparatively weak or negligible effects. Asterisks denote statistically significant effects.

**Figure S11:**
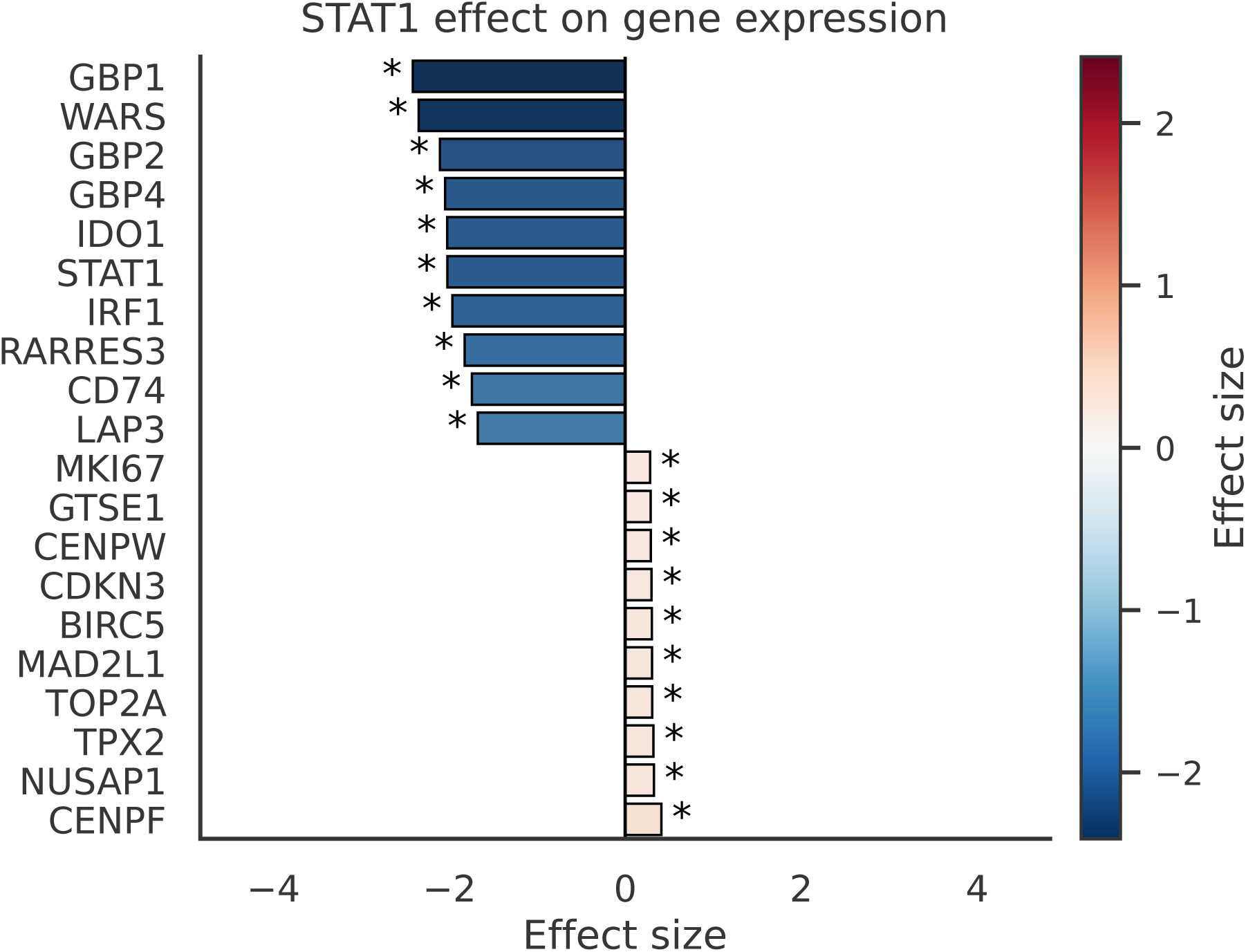
Top differential gene-expression effects associated with *STAT1* perturbation in the Perturb-CITE-seq melanoma dataset. Barplot showing the 10 strongest negative and 10 strongest positive inferred gene-expression effects for *STAT1* perturbation. Negative effects are enriched for canonical interferon-response genes, including *GBP1*, *WARS*, *GBP2*, *GBP4*, *IDO1*, *STAT1* and *IRF1*, consistent with loss of interferon signaling upon *STAT1* disruption. Positive effects include a smaller set of proliferation-associated genes, including *MKI67*, *TOP2A*, *TPX2* and *CENPF*. Asterisks denote statistically significant effects.

**Figure S12:**
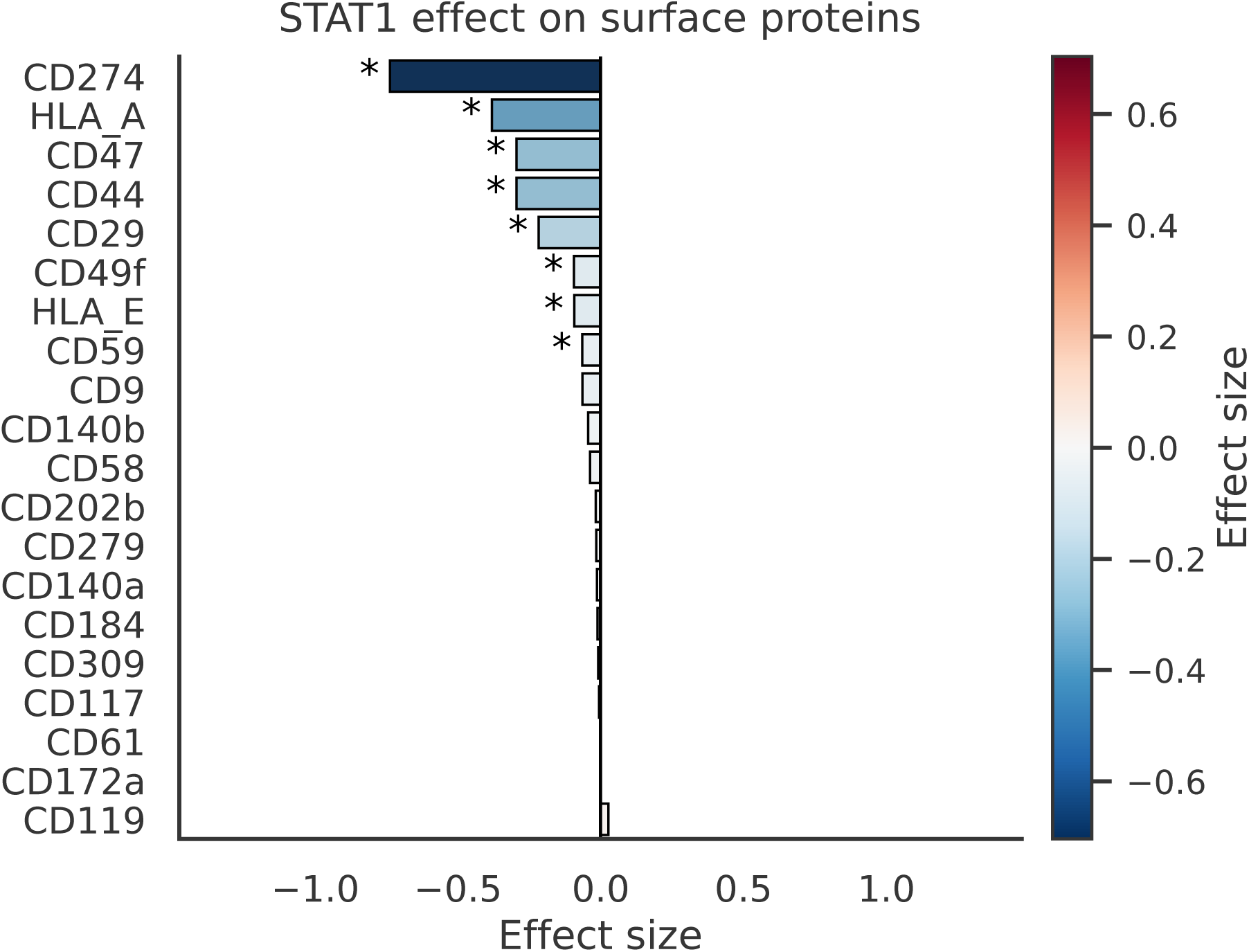
Differential surface-protein effects associated with *STAT1* perturbation in the PerturbCITE-seq melanoma dataset. Barplot showing inferred effects of *STAT1* perturbation on all measured surface proteins. Consistent with the pathway-level and gene-expression analyses, *STAT1* perturbation induces broad negative effects on immune-relevant surface markers, with the strongest suppression observed for *CD274* (PD-L1), *HLA A*, *CD47*, *CD44* and *CD29*. The remaining proteins show weak or negligible effects. Asterisks denote statistically significant effects.

**Figure S13:**
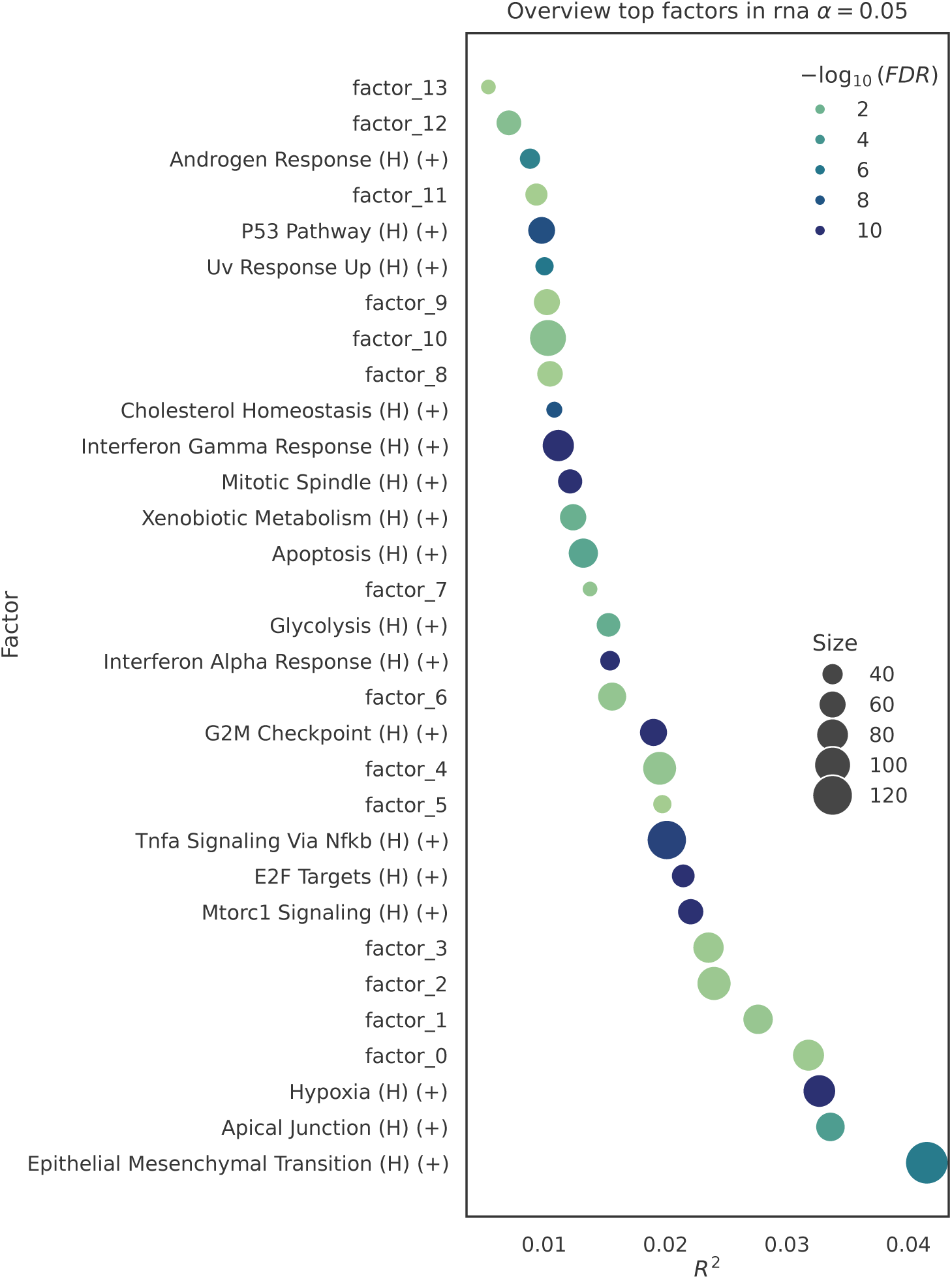
Overview of the latent factors in the Tahoe-100M perturbation atlas. Bubbleplot summarising the latent factors identified by PACMON at FDR < 0.05. The x-axis shows the fraction of variance explained (*R*^2^) by each factor, bubble size indicates factor size, and colour intensity corresponds to − log_10_(FDR) from principal component gene set enrichment (PCGSE). Pathway-informed factors recover major transcriptional programs including epithelial-mesenchymal transition, hypoxia, mTORC1 signaling, E2F targets, TNF*α* signaling via NF*κ*B, G2/M checkpoint and interferon-associated responses. Additional factors labeled as factor 0-factor 13 denote latent structure retained by the model but not significantly aligned with the available prior pathway annotations, consistent with the large scale and transcriptional diversity of the Tahoe-100M dataset relative to the limited number of informed Hallmark programs used here.

**Figure S14:**
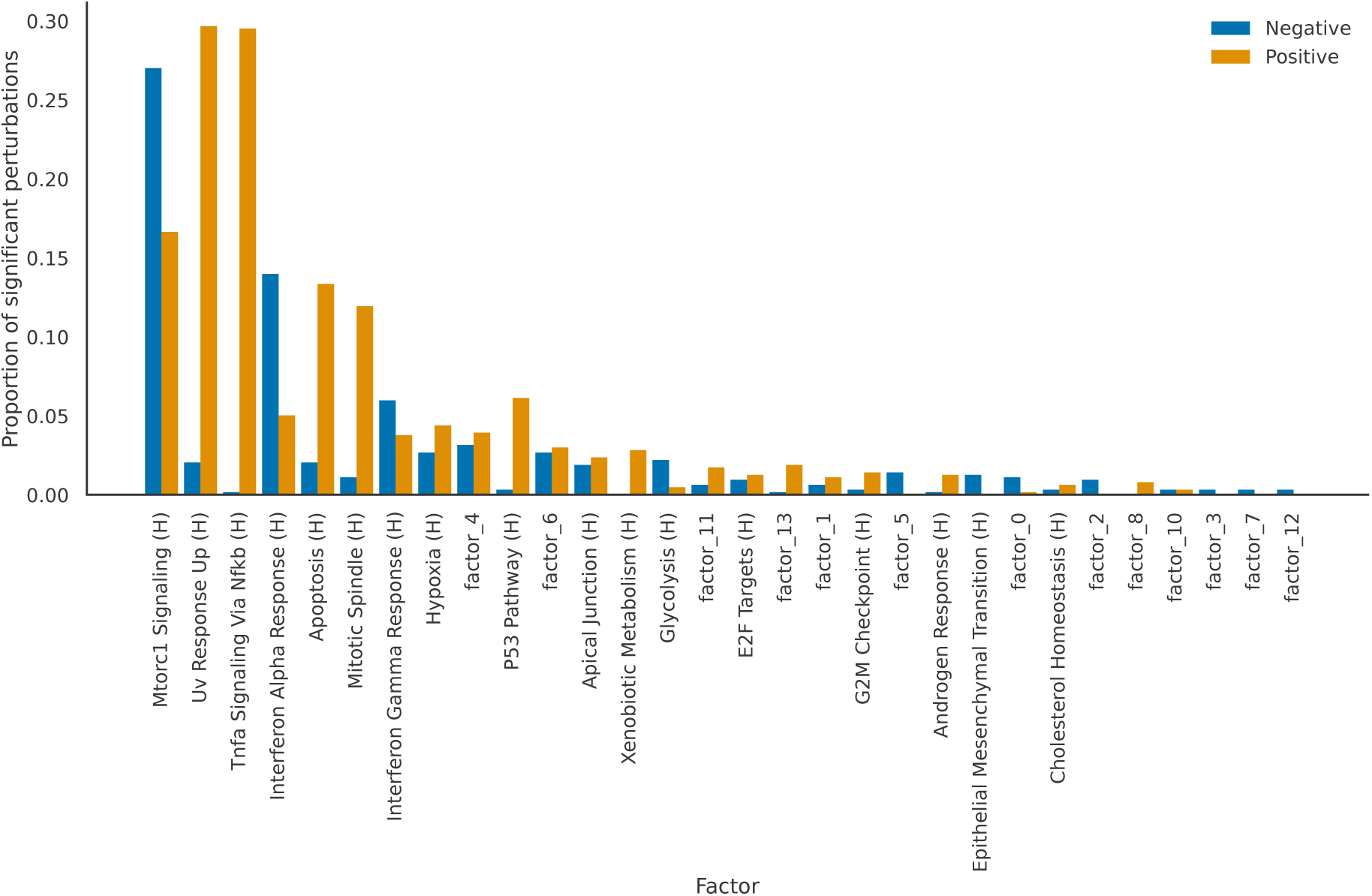
Sparsity of perturbation-pathway associations in the Tahoe-100M perturbation atlas. Paired barplot showing, for each latent factor, the proportion of drug-dosage perturbations with significant negative or positive associations. Factors are ordered by the total proportion of significant perturbations. Only a subset of pathway factors is recurrently affected across perturbations, and for each factor only a minority of perturbations show significant associations, consistent with a sparse perturbation-pathway structure. In particular, mTORC1 Signaling (H), UV Response Up (H), TNF*α* Signaling via NF*κ*B (H), interferon-associated programs, Apoptosis (H) and Hypoxia (H) account for the largest fractions of significant perturbations, whereas overwritten generic factors are affected less frequently and tend to appear toward the lower-ranked end of the distribution.

**Figure S15:**
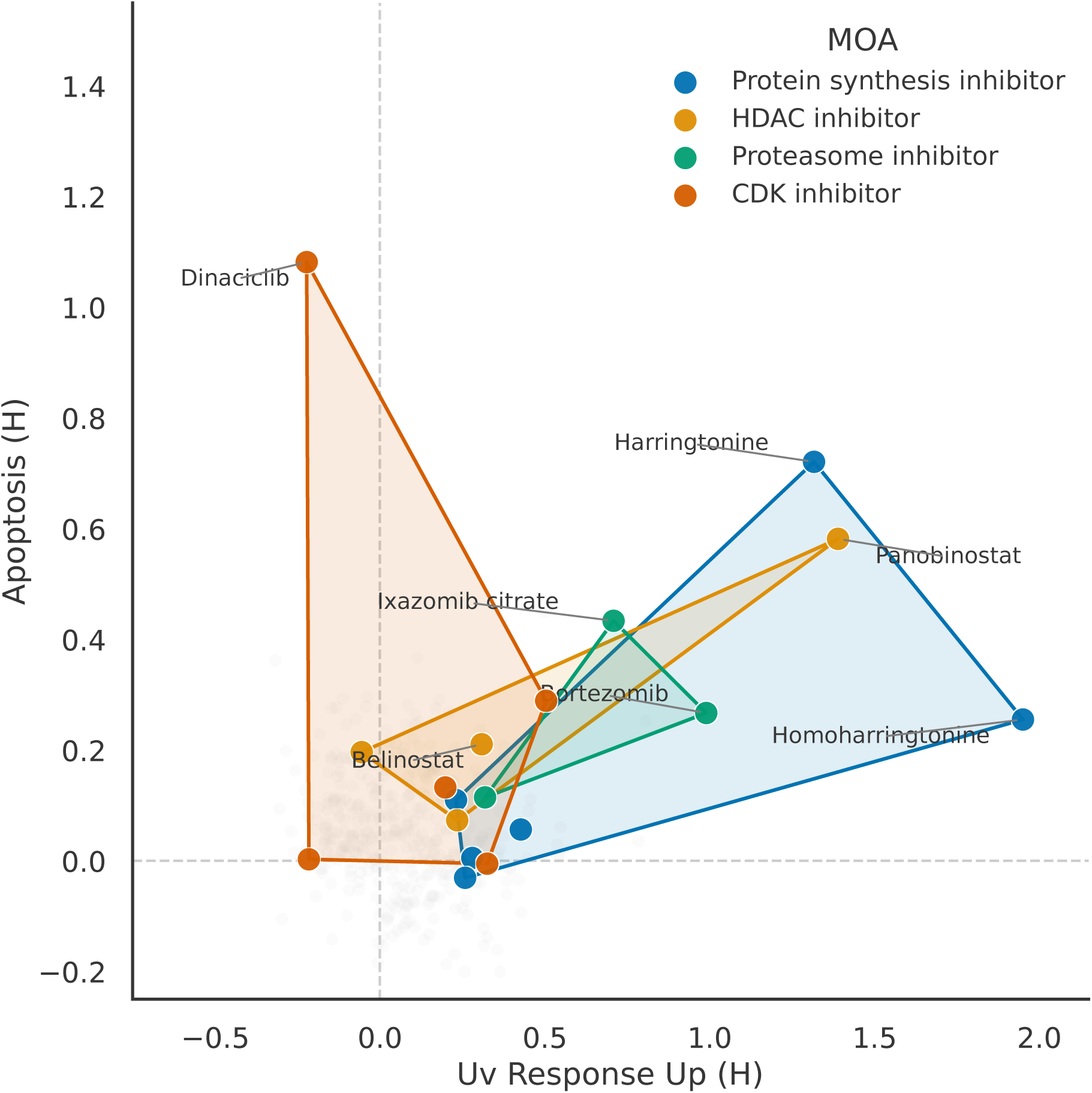
MOA-level projection of stress- and apoptosis-associated drug responses in the Tahoe100M perturbation atlas. Convex-hull projection of selected mechanisms of action (MOAs) in pathway space defined by UV Response Up (H) and Apoptosis (H). Each point corresponds to one drug after averaging pathway coefficients across the three assayed concentrations, and hulls summarise the distribution of drugs within each MOA class. Protein synthesis inhibitors occupy the strongest stress-associated region and extend toward high apoptosis, whereas HDAC, proteasome and CDK inhibitors show partially overlapping but distinct response profiles. Representative labeled drugs illustrate the heterogeneity of pathway responses within and across mechanisms of action.

**Figure S16:**
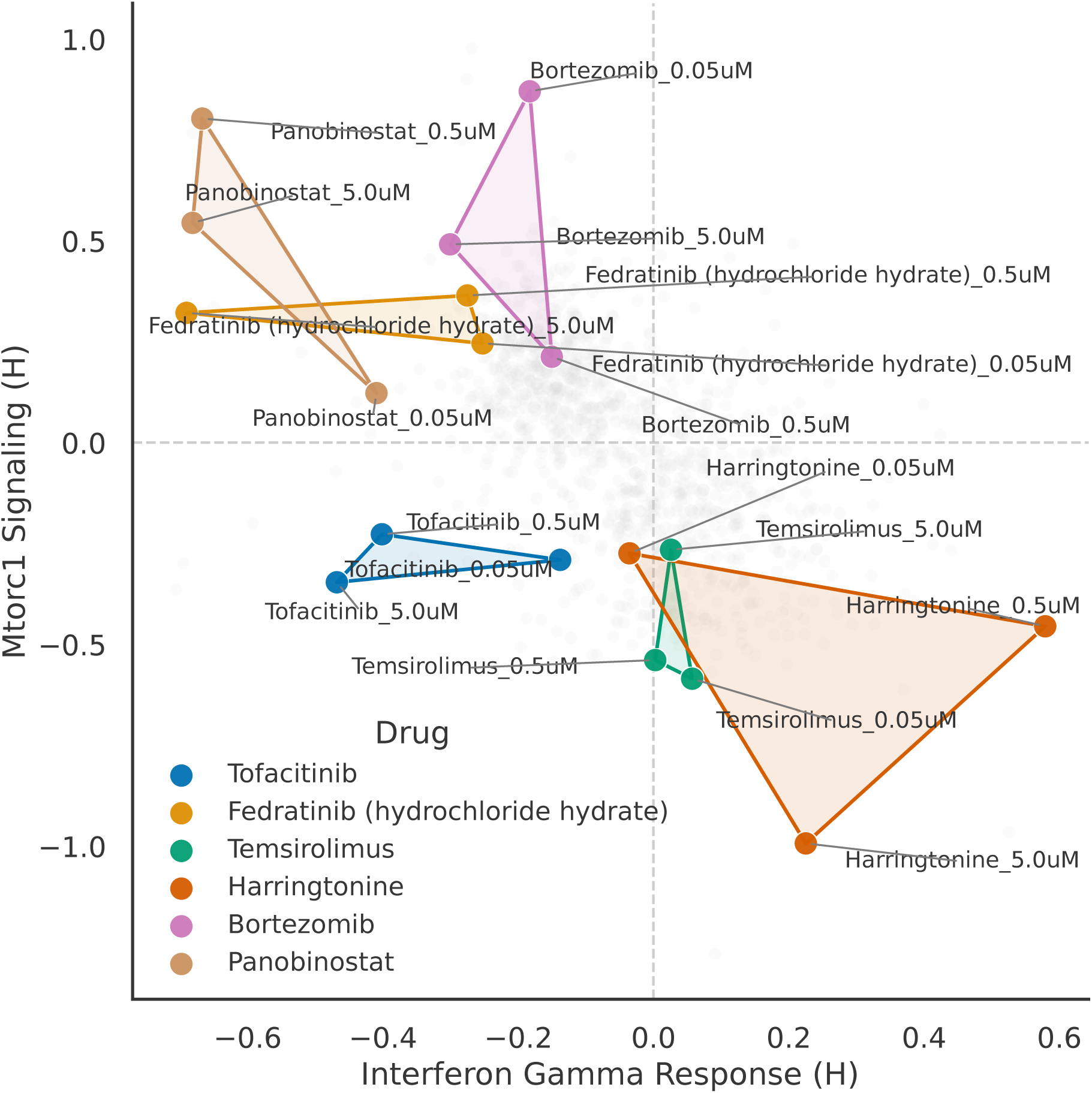
Drug-level projection of dose-dependent pathway responses in the Tahoe-100M perturbation atlas. Convex-hull projection of selected drugs in pathway space defined by Interferon Gamma Response (H) and mTORC1 Signaling (H). Each point corresponds to one assayed drug concentration, and hulls connect the dose-specific responses for each drug. This view provides a compact summary of how representative drugs traverse pathway space across increasing dosage while preserving the broader separation between interferon-associated and growth-associated response patterns.

**Figure S17:**
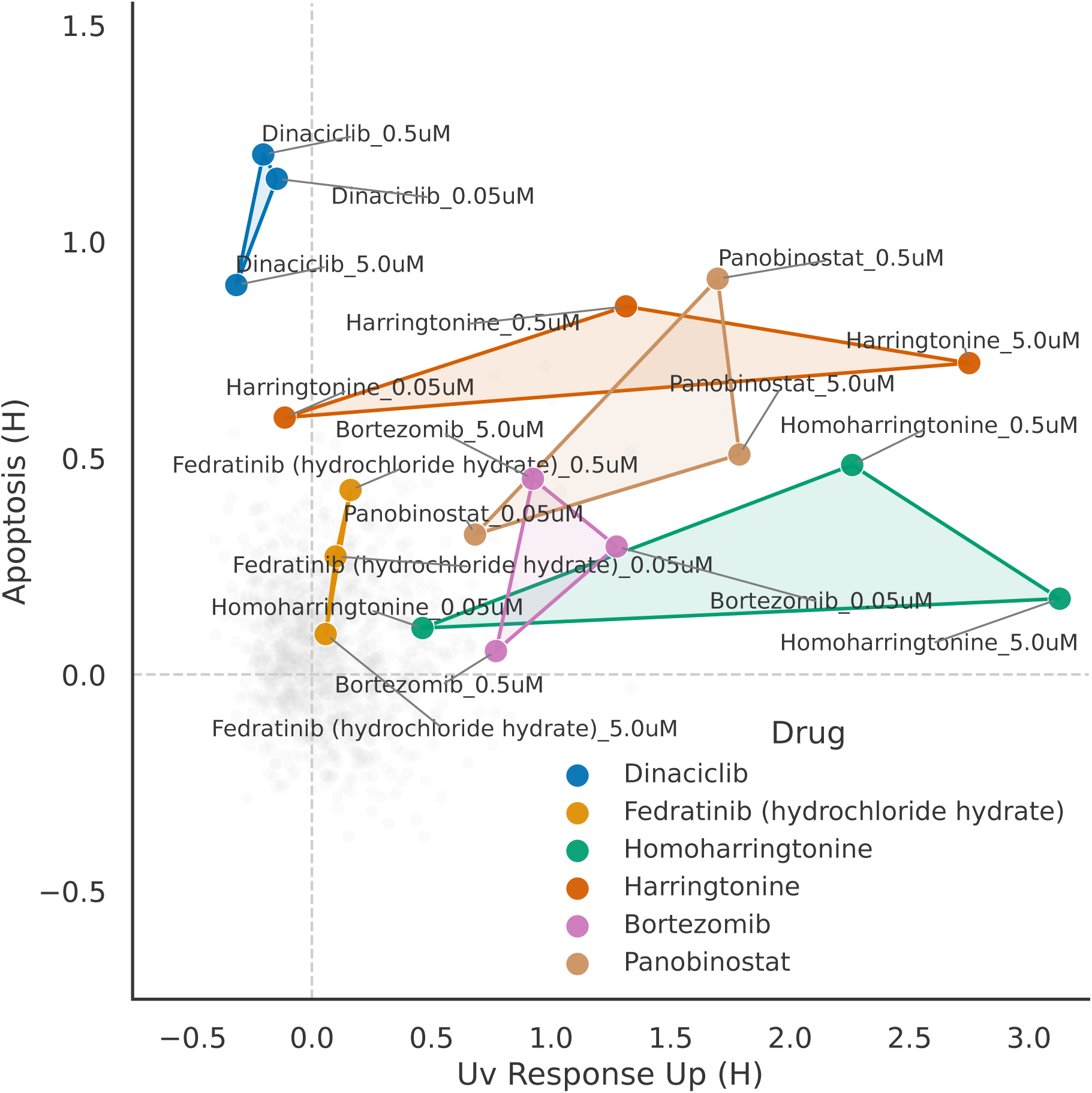
Drug-level projection of dose-dependent pathway responses in the Tahoe-100M perturbation atlas. Convex-hull projection of selected drugs in pathway space defined by UV Response Up (H) and Apoptosis (H). Each point corresponds to one assayed drug concentration, and hulls connect the dose-specific responses for each drug. This view summarises how representative drugs traverse pathway space across increasing dosage and highlights heterogeneity in dose-dependent responses across the selected compounds.

**Figure S18:**
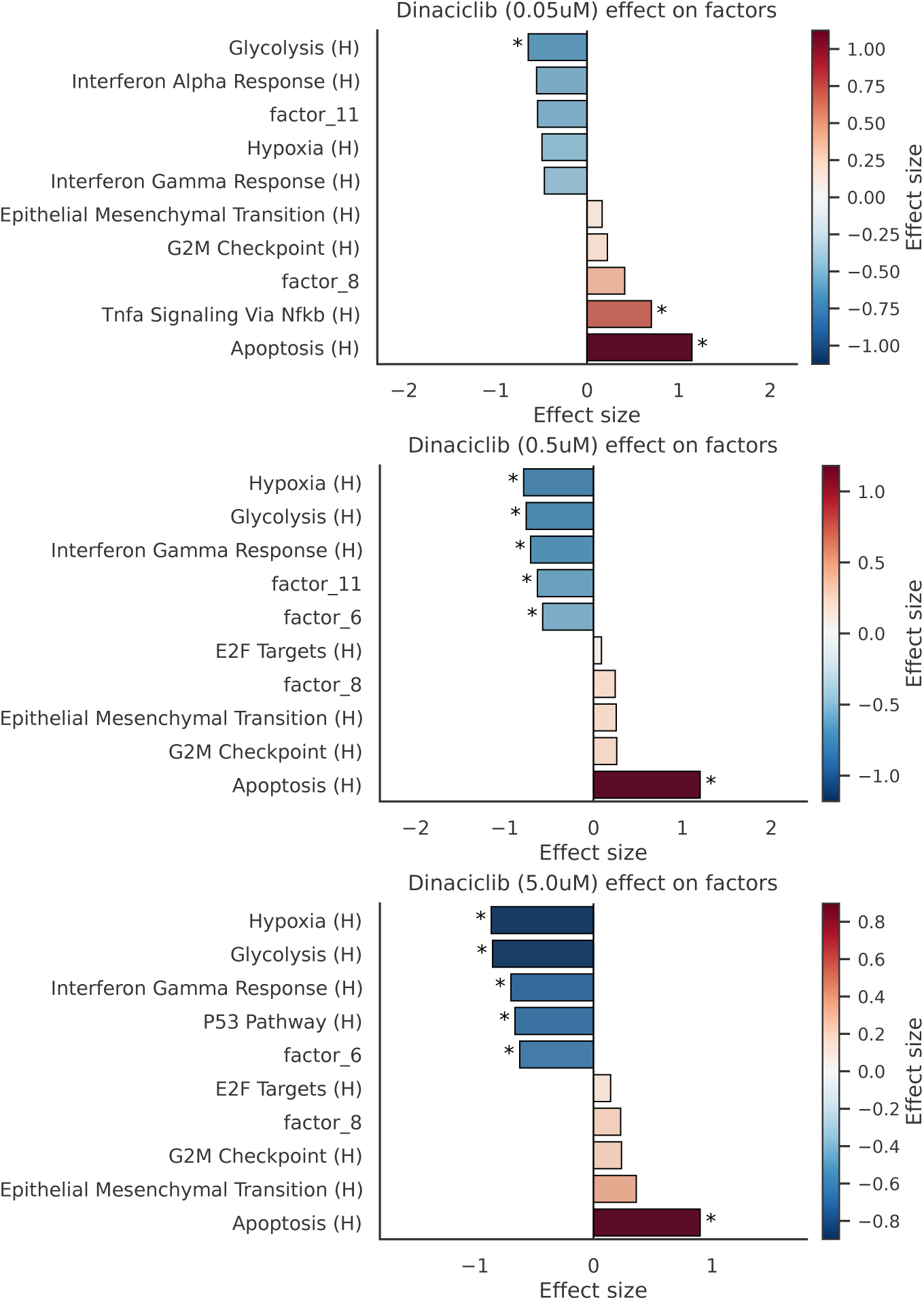
Dose-specific pathway effects of Dinaciclib in the Tahoe-100M perturbation atlas. Barplots showing the strongest inferred pathway effects for Dinaciclib at 0.05 *µ*M (top), 0.5 *µ*M (middle) and 5.0 *µ*M (bottom). Negative effect sizes indicate pathway suppression, whereas positive values indicate increased pathway activity. Across all three concentrations, Dinaciclib is consistently associated with increased apoptosis and reduced activity of interferon- and metabolism-associated programs, while additional pathway effects vary with dosage. Asterisks denote statistically significant effects.

**Figure S19:**
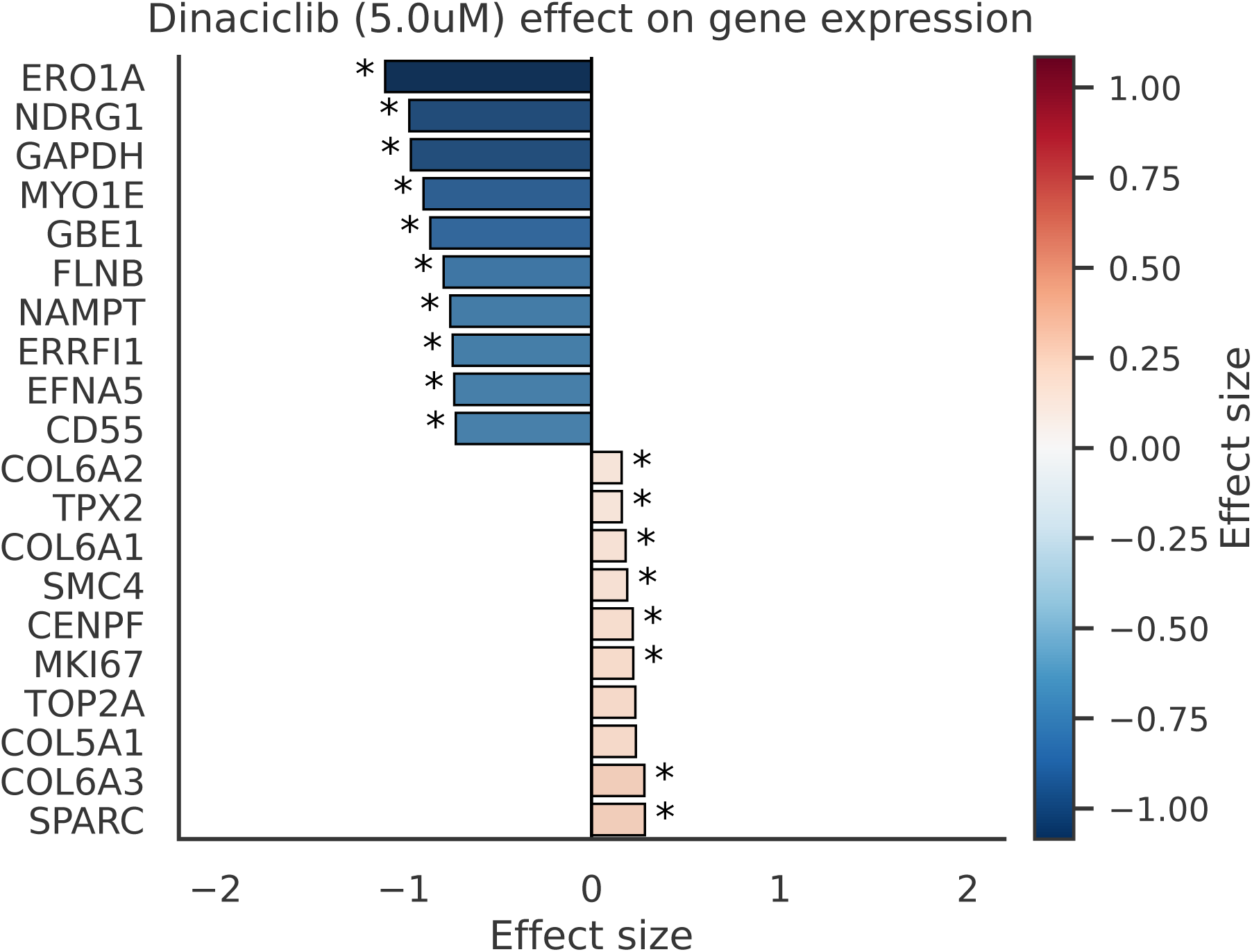
Top differential gene-expression effects associated with Dinaciclib at 5.0 *µ*M in the Tahoe100M perturbation atlas. Barplot showing the 10 strongest negative and 10 strongest positive inferred gene-expression effects for Dinaciclib at the highest assayed concentration. Negative effects are enriched for genes associated with hypoxia and metabolic adaptation, whereas positive effects include a smaller set of genes related to proliferation and extracellular-matrix organization.

**Figure S20:**
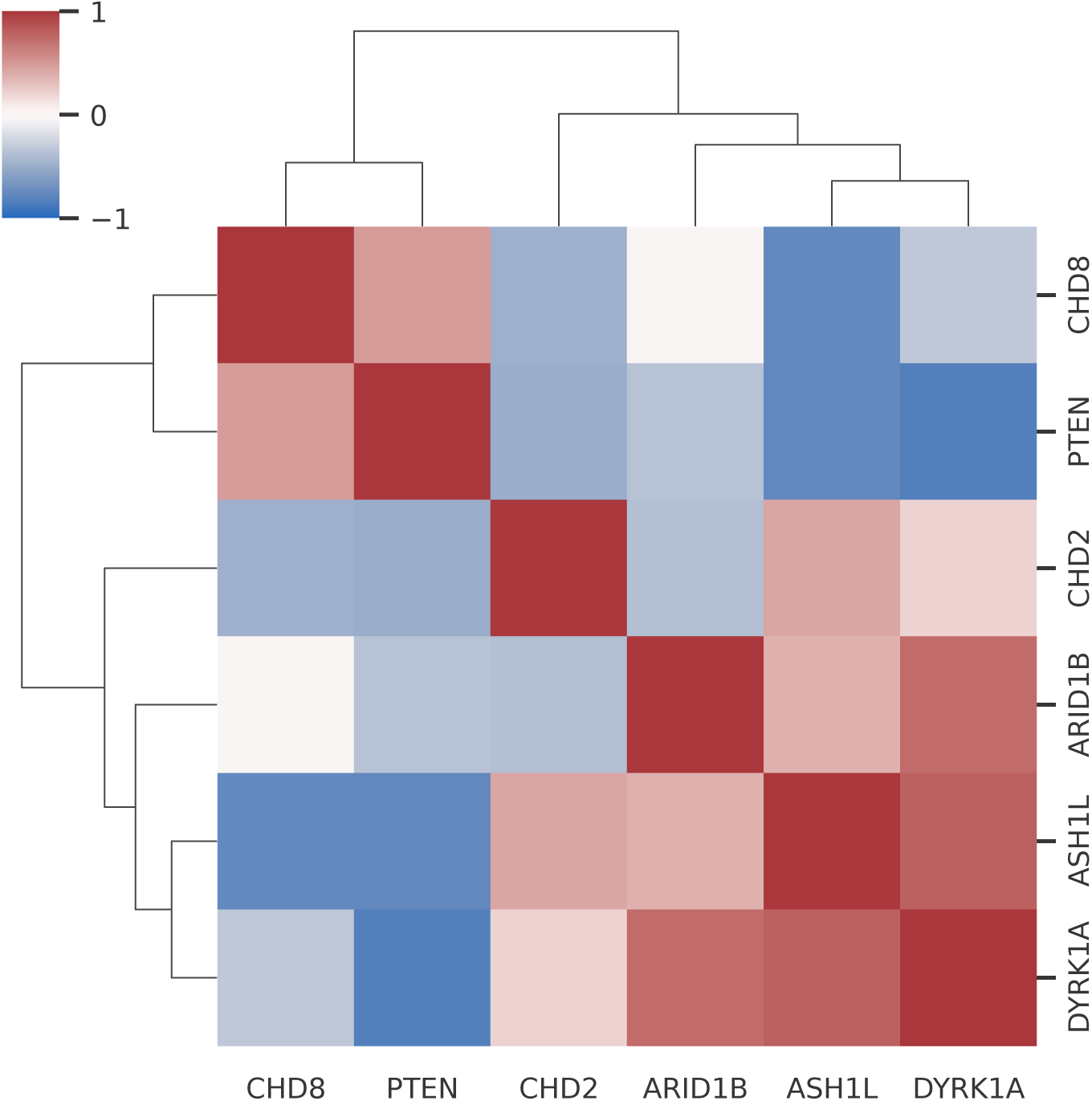
Heatmap visualising the cosine similarity computed for the perturbation-factor coefficient vectors of selected perturbations from the LUHMES dataset. Factors explaining 98% of the variance were used for the computation. The similarity structure reveals two clusters: CHD8 and PTEN have very similar perturbation effects, whereas CHD2, ARID1B, ASH1L, and DYRK1A form a second cluster.

**Figure S21:**
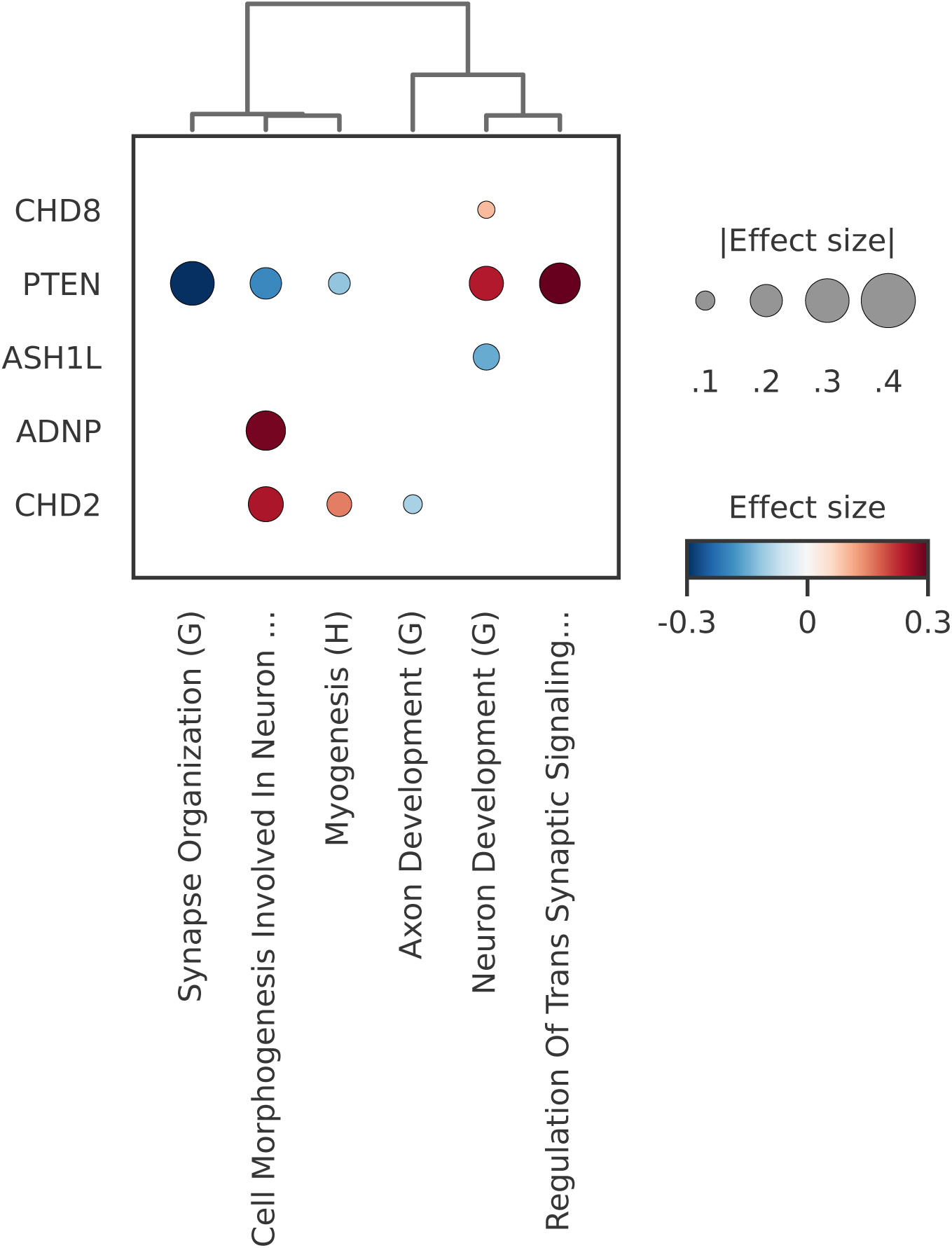
Perturbation-factor coefficients for the LUHMES dataset. ”Cell Morphogenesis Involved In Neuron …” denotes the GOBP pathway ”Cell Morphogenesis Involved In Neuron Differentiation (G)”. ”Regulation of Trans Synaptic Signaling…” is based on the corresponding GOBP pathway. Dot colour denotes effect direction and magnitude, and dot size denotes absolute effect size. Clustering the factors by their associated perturbations reveals two clusters, which are mainly driven by the *PTEN* perturbation effect.

**Figure S22:**
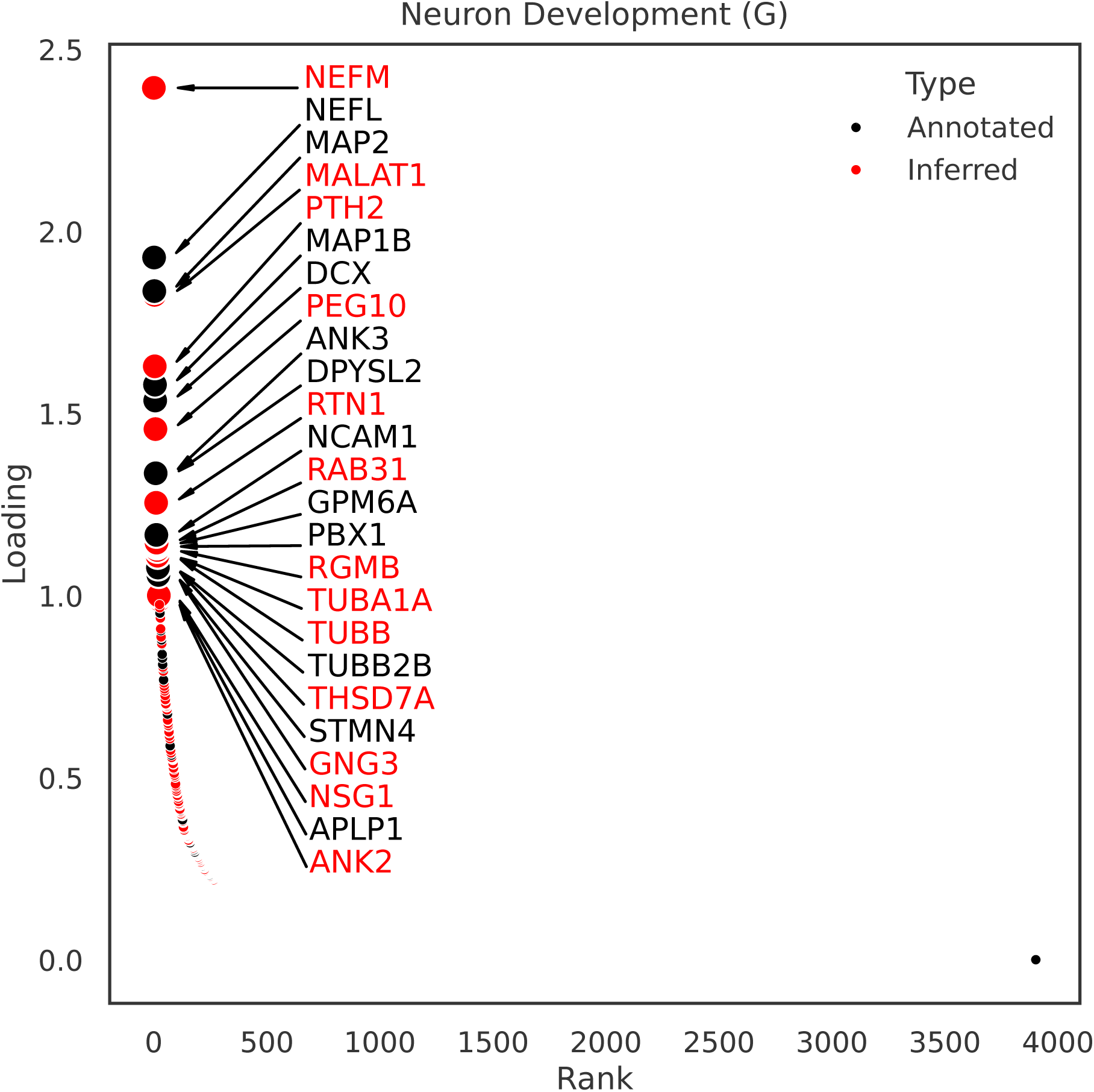
Factor loadings for one exemplary LUHMES factor related to neuron development. The pathway definition is based on the GOBP gene set. Genes are ranked by their individual weight on the factor score. Black genes are annotated in the prior GOBP pathway definition, while red genes are inferred by the model.

## 7 Supplementary Tables

**Supplementary Table S1.** Complete perturbation-pathway association matrix for the PerturbCITE-seq analysis. Provided as a CSV file.

**Supplementary Table S2.** Complete perturbation-gene effect matrix for the Perturb-CITE-seq analysis. Provided as a CSV file.

**Supplementary Table S3.** Complete perturbation-protein abundance effect matrix for the Perturb-CITE-seq analysis. Provided as a CSV file.

**Supplementary Table S4.** Complete perturbation-pathway association matrix for the Tahoe-100M analysis. Provided as a CSV file.

**Supplementary Table S5.** Complete perturbation-gene effect matrix for the Tahoe-100M analysis. Provided as a CSV file.

